# Functional maps of the primate cortex revealed by through-skull wide-field optical imaging

**DOI:** 10.1101/2020.12.05.413047

**Authors:** Xindong Song, Yueqi Guo, Hongbo Li, Chenggang Chen, Zachary Schmidt, Xiaoqin Wang

## Abstract

The primate cerebral cortex is organized into specialized areas representing different functional modalities (e.g., vision, audition, touch) and their associations along a continuous surface. The functional maps of these areas, however, are often investigated in a single modality at a time. Here, we developed and applied to awake primates a polarization-enhanced wide-field optical imaging method for measuring cortical hemodynamics through the intact skull. Adjacent somatosensory, auditory, and visual cortices were noninvasively localized and rapidly parcellated in awake marmosets (*Callithrix jacchus*), a primate model featuring a smooth cortex. Detailed somatotopy, tonotopy, and retinotopy were also mapped out on an individual-subject basis, with a new pure-tone-responsive tonotopic gradient discovered outside the auditory core. Moreover, the motion-sensitive extent surrounding the primate-specific MT/V5 and the location of a face-sensitive patch were both revealed with respect to retinotopy. This approach provides a powerful tool for mapping the functional landscape across modalities in a single non-human primate subject, and thus opens new opportunities for probing how primate cortical system is organized to enable real-world behaviors.

## INTRODUCTION

Evolution has endowed primates with unique features in the organizations of cerebral cortical areas that contribute to our remarkable mental abilities^1^. Information from various sensory modalities is routed through these different areas that are located along a continuum of cortical surface. The functional organizations of the cortical areas, however, are usually studied one area or one modality at a time. Our ability to efficiently map primate cortex on its continuous surface across cortical areas and modalities is still limited. Enabling such an ability would thus lay a foundation for further systematic investigations into how different parts of the primate cortical system are organized together to generate our real-world perceptions and behaviors. Here we demonstrate a through-skull wide-field optical imaging strategy to enable observations across somatosensory, auditory, and visual cortices in individual awake marmoset monkeys.

Wide-field optical imaging across multiple modalities in the cortex, although has recently been applied through-skull in awake mice^2, 3^, has so far been impractical in primates. The imaging view and thus the modality coverage are constrained by the practical size of skull and dura openings, which were considered necessary in primates^4, 5^. Across primate species, both skull thickness^6^ and the extent of cortical folding^7^ vary substantially (Fig. 1a). Interestingly, the common marmoset (*Callithrix jacchus*), an increasingly popular biomedical and neuroscientific primate model^8^, features a skull thickness of ∼0.5mm (6) and a cortical folding index of ∼1.2 ^(7)^. Both numbers are among the lowest of commonly used non-human primate (NHP) models (Fig. 1a), indicating a possibility of imaging through the intact skull and resolving continuous topographic organizations in the cortex. Since the marmoset cortex is mostly smooth without folding (except for the lateral sulcus), many cortical areas are directly accessible under the skull. Noticeably, the middle temporal visual field (MT/V5)^9^, most of the auditory cortex^10^, and the lateral part of the primary somatosensory area 3b^11^ are all located near but outside of the lateral sulcus (Fig. 1b) and together provide an interesting target to image across modalities. Therefore, we designed a chronic head cap implant with a lateral recording chamber (Fig. 1c), the bottom of which consists of a thin layer of dental cement directly covering the un-thinned intact skull. As activity increases in a cortical region under the skull, a hemodynamic response follows, which in turn changes the cortical tissue’s optical properties, including absorption^12, 13^. If both skull and dura are surgically removed and the brain is illuminated, this change can be picked up by the intensity of backscattered light and imaged by a camera (referred to as intrinsic optical signal imaging) in a variety of species including NHPs^4, 5, 12–15^ (in one case^16^, through locally thinned skull). The same imaging approach applied through the intact skull without thinning, although has been feasible in mice^17–20^, is technically challenging in other species such as marmosets. Due to the thickness of marmoset skull, the prominent optical scattering in the bone (Extended Data Fig. 1a) causes many photons to be backscattered before even reaching the cortex and reduces the imaging sensitivity to map cortical functions. Furthermore, as the skull curvature varies across the recording chamber, it is difficult to illuminate a large area homogenously without saturating pixels from surface reflection (Fig. 1d). To address these problems, we designed a cross-linear polarization enhanced wide-field imaging setup (termed XINTRINSIC, Extended Data Fig. 1c). By illuminating the skull surface with linearly polarized light and recording backscattered light only of the orthogonal (cross-linear) polarization, the surface reflection was effectively eliminated, and the entire chamber was homogeneously illuminated (Fig. 1e). Moreover, since only light that undergoes multiple scattering events is significantly depolarized and therefore received by the camera, this strategy rejects the photons that only visit shallower structures^21^ (Extended Data Fig. 1d). After testing a broad range of wavelengths (470-850nm), we found that the shorter wavelengths (e.g. blue and green) can better detect hemodynamics through the marmoset skull (Extended Data Fig. 2). Since our green light (530 nm) is near an isosbestic point, at which the change in total hemoglobin concentration can be measured independently from the change in blood oxygenation^13^ (Extended Data Fig. 1b), we sought to map cortical functions through the intact marmoset skull with green light. The lateral resolution (full width at half maximum [FWHM]) of XINTRINSIC under this condition was estimated as 0.63mm (Extended Data Fig. 1e).

**Fig. 1.**
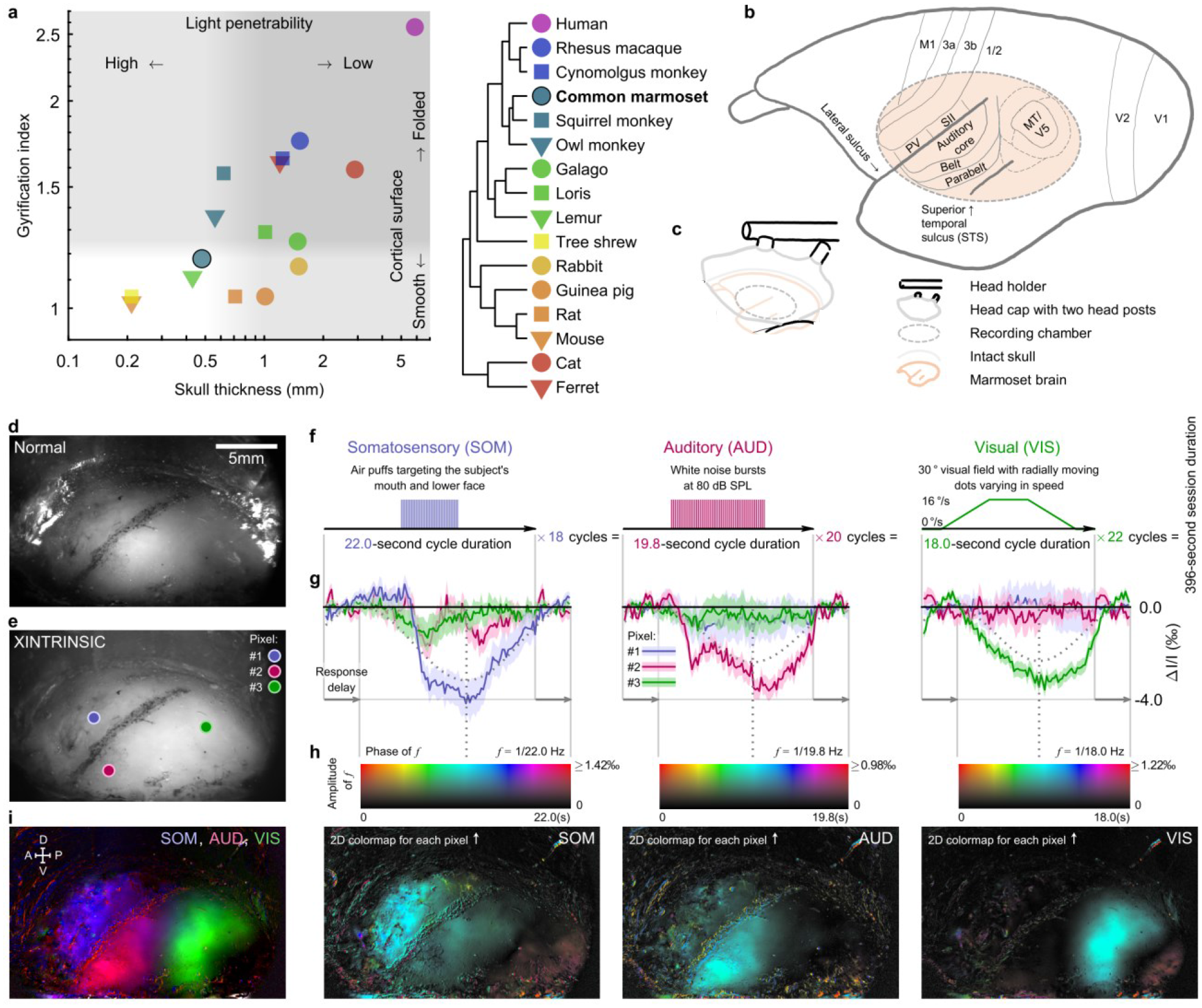
Marmoset through-skull imaging and functional parcellation of cortical modalities. **a**, Gyrification index^7^ and skull thickness^6^ are compared across species (more details on data sources in Methods). Light penetrability decreases as the skull thickness increases. Increased cortical folding (indicated by larger gyrification index) fragmentizes functional maps in the imaging field. **b**, A sketch of key cortical areas in marmosets^9–11^. **c**, Illustration of a head-cap implant and recording chamber. **d, e**, A view of the recording chamber imaged with (**e**) or without (**d**) XINTRINSIC enhancement using green light. The putative lateral sulcus was marked during implant surgery. **f**, Stimuli design for functionally parcellating somatosensory (SOM), auditory (AUD), and visual (VIS) cortices. **g**, Intrinsic signal response to the stimuli shown in **f** from three exemplar pixels, whose locations are labeled in **e** with the corresponding colors of the signal traces. Each average response trace is shown as a solid line with a shade representing the errors (standard error mean, SEM). Stimulus repetitions in each modality: 18 for SOM (left), 20 for AUD (middle), 22 for VIS (right). For each modality, a cosine “fitting” curve to the trace of the most responsive exemplar pixel was drawn (dotted cosine line) with the response phase indicated by a dotted vertical line. **h**, Response maps of three modalities (left: SOM, middle: AUD, right: VIS). For each pixel, the cosine “fitting” phase and amplitude were encoded respectively as color and intensity on the map, according to the 2D colormap shown above each modality map. **i**, A summary map of functional modality parcellation, with each modality represented by a color (blue: SOM, red: AUD, green: VIS). A (anterior), P (posterior), D (dorsal), V (ventral).

## RESULTS

### Functional modality parcellation of marmoset cortex through-skull

To functionally parcellate somatosensory, auditory, and visual cortices, we designed an experiment in which periodic stimuli of each modality were locked to distinct repetition periods within the same session (Fig. 1f). The somatosensory stimuli were mild air-puffs delivered to the subject’s mouth and lower face presented at the middle of each 22-second-long cycle (Fig. 1f left). The auditory stimuli were 80 dB SPL white noise bursts presented at the middle of each 19.8-second-long cycle (Fig. 1f middle). The visual stimuli were radially outward moving dots gradually speeding up and then slowing down within each 18-second-long cycle (Fig. 1f right). Within a 396-second-long session, somatosensory, auditory, and visual stimuli were repeated for 18, 20, and 22 cycles, respectively. The spectrum of each pixel’s response was calculated across the entire session with compensations for estimated delay and polarity. If the signal of a pixel followed a particular type of stimuli well, it would show an evoked amplitude at the corresponding frequency in the spectrum with a phase near the middle of the stimulus cycle^18, 22^ (examples shown in Fig. 1g). For every type of stimuli, we plotted each pixel’s response by showing its phase at the corresponding frequency with color and the amplitude of that frequency with intensity (from traces as those shown in Fig. 1g to colormaps in Fig. 1h). Each type of sensory stimuli evoked responses in a distinct region within the imaging chamber. The orofacial air-puffs activated the cortical region anterodorsal to the putative lateral sulcus (Fig. 1h left), whereas the white noise sound activated the region posteroventral to it (Fig. 1h center). Additionally, moving dots activated the region posterior to the auditory region (Fig. 1h right). These locations are consistent with previous results using invasive electrophysiology methods^11, 23, 24^. Interestingly, there was a low-intensity gap between the auditory and visual regions that cannot be effectively activated by any of the current stimuli (Fig. 1i). A large part of this location is consistent with regions suggested for audiovisual integration of motion in macaques and humans^25^. Although the general phenomena described above were robust across all tested subjects, individual variance was also evident (Extended Data Fig. 3), which demonstrates the importance of parcellating these modalities on an individual-subject basis. Data from this experiment thus showed the ability of XINTRINSIC to quickly map such a functional landscape consisting of multiple modalities.

### Through-skull somatotopy mapping

After functionally parcellating these modalities, we sought to map topographic organizations within each individual modality in the same subject, starting with somatotopy. The primary somatosensory area 3b in marmosets, similar to other primates, has a lateral-to-medial representation of the contralateral body surface (or musculature) from head to foot^11, 26^. Moreover, part of the secondary somatosensory area (SII) and parietal ventral area (PV) are also exposed outside the lateral sulcus^11^ (Fig. 2a). Electrophysiology mapping in anesthetized marmosets suggests that area 3b shares an orofacial representing border with both SII and PV^11^, consistent with the orofacial region located in our experiment (Fig. 1h left). However, it is less clear whether the representations of the mouth and the face are differentiable topographically. To test this in another experiment, we moved one air-puff nozzle closer to the teeth to target the oral cavity and added another air-puff nozzle to target the contralateral cheek (Fig. 2b). These two nozzles delivered air-puffs at different cycles within the same session (Fig. 2c). While both nozzles evoked cortical activations in regions immediately anterodorsal to the putative lateral sulcus (Fig. 2d), the oral region appeared more anteroventral to the major facial region (Fig. 2e, f), with minimal overlap between them in all tested subjects (Extended Data Fig. 4), suggesting that these two cortical representations are topographically differentiable and together form an oral-facial somatotopic gradient in marmosets.

**Fig. 2.**
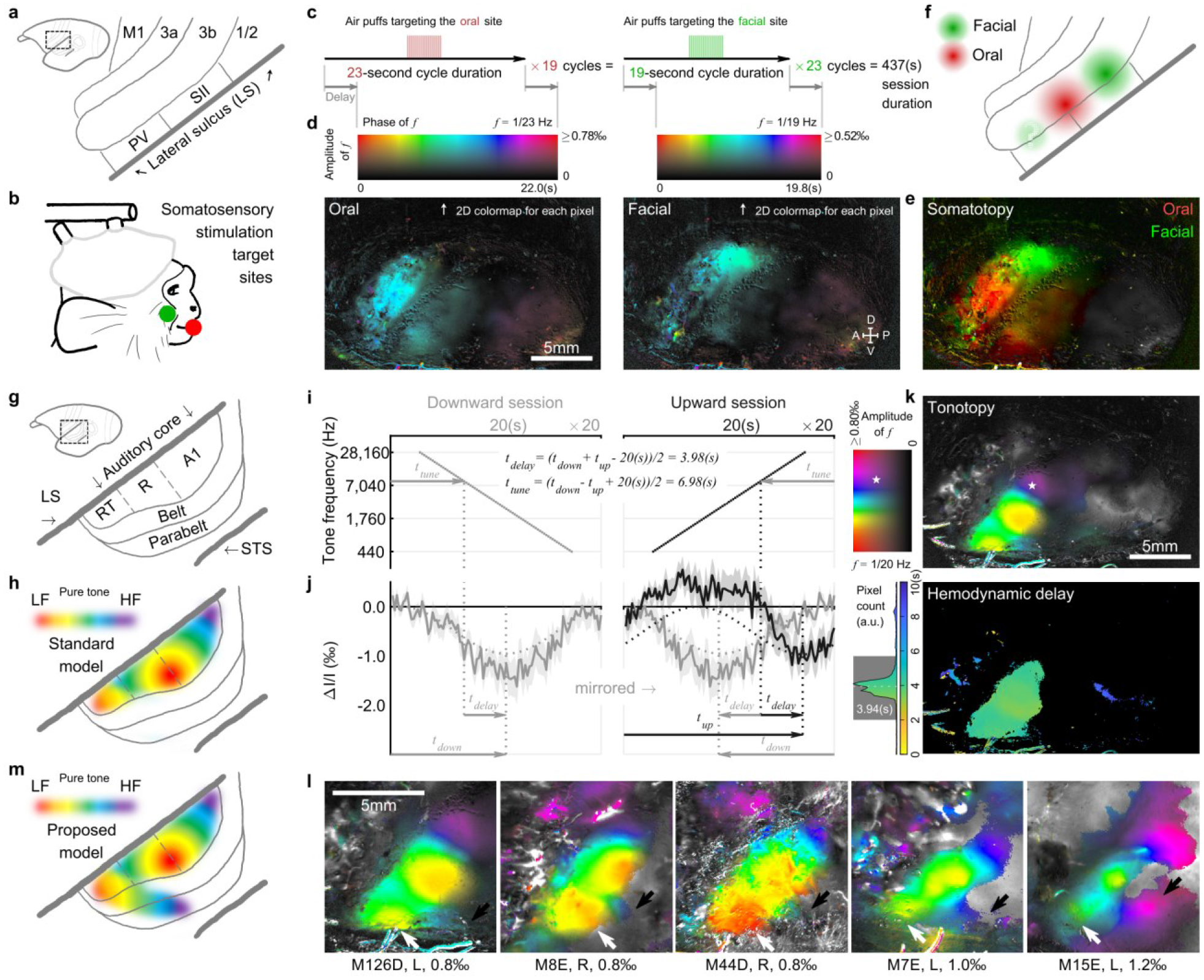
Through-skull somatotopy and tonotopy mapping. **a**, A sketch of marmoset somatosensory cortices^11^. **b**, The target sites for air-puff stimulation. The same colors (oral: red, facial: green) were used in **c**, **e**, and **f** as well. **c**, Stimuli design for mapping oral and facial representations in somatosensory cortices. **d**, Response maps evoked by oral (left) or facial (right) stimuli. For each pixel, the response phase and amplitude are visualized respectively as color and intensity on the map. **e**, A summary response map of the orofacial somatotopic gradient. Oral (red) and facial (green) representations are well differentiable. **f**, An illustration of oral and facial response areas in somatosensory cortices (estimated). **g**, A sketch of marmoset auditory cortices^10^. **h**, The standard model of pure-tone-responsive tonotopic gradients in marmoset auditory cortices^23, 27^, LF: low frequency, HF: high frequency. **i**, Stimuli design for mapping tonotopy using a downward (left) or an upward (right) frequency sequence in each cycle. **j**, Intrinsic signal (solid lines: average traces, shades: SEMs, n=20) responding to the stimuli shown in **i** from an exemplar pixel (location labeled by a pentagram in **k**). Dotted lines: response phase extraction (same as in Fig. 1g). Arrows and equations: tone tuning and hemodynamic delay estimation process. **k**, Tonotopy (top) and hemodynamic delay map (bottom). The colormap (top left) shows 2D codes for visualizing each pixel’s tone-tuning frequency (color) and response amplitude (intensity). The hemodynamic delay values are color-coded (for top 10% of most responsive pixels) and counted in the histogram (bottom left). Rectangular gray shade: the acceptable delay range. The average delay within this range is 3.94 seconds. **l**, Tonotopy from five tested subjects. White (black) arrows: low (high) frequencies of the newly discovered tonotopic gradient. Subject ID, hemisphere (L or R), and saturated display amplitude are listed below each plot. Right hemispheres are mirrored to the left for display purposes. **m**, Proposed model for pure-tone-responsive tonotopic gradients in marmoset auditory cortex. A newly discovered pure-tone-responsive tonotopic gradient extends from the rostrotemporal field (RT) of the auditory core to the putative parabelt region.

### Through-skull tonotopy mapping

Looking across the lateral sulcus, we sought to localize tonotopic maps in the auditory cortex using pure tones. The primate auditory cortex exhibits a hierarchical progression from the core region to belt and parabelt regions^10, 27^ (Fig. 2g). The core region is suggested to have three primary-like areas: A1, R (rostral field), and RT (rostrotemporal field) in a caudal to rostral order (Fig. 2g), all of which are responsive to pure tones^23, 27^ (Fig. 2h). Both A1 and R are clearly tonotopically organized and share a low-frequency reversal in between^23, 27^ (Fig. 2h). Although the tonotopy in RT is less clear due to limited sampling in previous studies, a high-frequency reversal has generally been used to identify the transition from R to RT^15, 23, 27–29^. To map pure-tone-responsive tonotopic gradients, we utilized a phase-encoded Fourier approach^18, 30^. A pure-tone pip sequence increasing or decreasing in frequency by discrete semitone steps was played at a moderate sound level (50 dB SPL) for 20 cycles in each recording session (Fig. 2i). The responses from these two sessions (Fig. 2j) were then combined to derive each pixel’s tone tuning (Fig. 2k upper) and hemodynamic delay (Fig. 2k lower). The tone tuning was visualized with color (red: low-frequency, purple: high-frequency), while the tone responsiveness was visualized as the intensity in the map (bright: responsive, dark: non-responsive). Clear tone responsiveness and tonotopic gradients were found largely within the region responsive to noise stimuli (Fig. 1h center) and abutted the putative lateral sulcus (Fig. 2k upper). A zoomed-in view is shown in Fig. 2l (leftmost) together with those of all other tested subjects (also see Extended Data Fig. 5). The known tonotopic gradients of A1-R-RT (Fig. 2H) were evident in every subject. In addition, we observed a new low-to-high tonotopic gradient extending laterally and caudally from the low-frequency part of RT (white arrows in Fig. 2L) towards a high-frequency region outside the core region (black arrows in Fig. 2L). These gradients were further confirmed by direct imaging of the cortex through a subsequently implanted cranial window in one of the subjects (Extended Data Fig. 6). The existence of a pure-tone-responsive tonotopic gradient beyond the classical tonotopic maps calls for a revision of our view on the tonotopic organizations of primate auditory cortex (Fig. 2m).

### Through-skull retinotopy and motion sensitivity mapping in MT/V5 complex

Next, we sought to map retinotopic gradients of the moving-dots-sensitive visual region determined in the parcellation experiment. This region presumably includes MT/V5^(31)^ (Fig. 3a), a motion-sensitive visual area^24, 32, 33^ believed to be shared by all primates^1^. In fact, most neurons within MT/V5 are clearly selective to motion direction, largely regardless of other visual features like form^9, 24, 32^. What is less clear, is to what degree the moving-dots-sensitive region includes areas adjacent to MT/V5, since a complex of satellite areas surrounding MT/V5 is largely considered to be involved in motion processing as well^9, 24, 33, 34^ (Fig. 3a). Nonetheless, some features regarding retinotopic organization are generally consistent across studies in several species including marmosets (Fig. 3b): (i) MT/V5 is retinotopically organized^24, 31, 33–36^ to represent the entire contralateral visual field in a non-mirrored image fashion^37^ and shares a common foveal representation at its ventral side with its neighboring areas; (ii) The region anterior to MT/V5 has a crude retinotopic organization but as a mirrored image with a reversed polar angle gradient^24, 35, 36^. Yet, the spatial relationship between these retinotopic features and moving-dots sensitivity in the marmoset MT/V5 complex is not clear. Here, we first confirmed the spatial extent of the motion sensitivity to moving dots was robust and replicable, by repeating the visual stimuli used in the parcellation experiment without the stimuli of any other modality. (Fig. 3c, d vs Fig. 1f, h right). Next, to map retinotopic eccentricity, we varied the range of moving dot stimuli between the center and periphery of the visual field (Fig. 3f). As a result, a periphery-preferring region was located at the anterodorsal part of the motion-sensitive region, while a center-preferring region occupied the posteroventral side of the motion-sensitive region (Fig. 3g). Moreover, to map retinotopic polar angle, the dots were permitted to move radially only in a quarter-circle shaped range in the visual field that was swept either clockwise (Fig. 3i) or counterclockwise (Fig. 3l) for 20 rounds. Similar to the tonotopy experiment, polar angle tuning and hemodynamic delay were calculated by combining these “reversed” sessions together (Fig. 3j, m, o). The resulting lower-upper-lower polar angle gradients for representing the contralateral field were virtually orthogonal to the eccentricity gradient (Fig. 3o vs Fig. 3g). Putting these together, the extent of moving-dots sensitivity occupied two retinotopic regions, and each had a representation of the entire contralateral visual field. The more posterodorsal, non-mirrored retinotopic region was consistent with MT/V5, and the more anteroventral region was retinotopically organized in a mirrored image. These observations were generally consistent in all subjects we tested (Extended Data Fig. 7). It is worth noting that we did not train our subjects to fixate during these sessions. Nevertheless, calibrated eye-tracking results showed the subject did actively gaze to the center, with the median gaze position of any session maximally 1.1° off the center, while the motion stimuli subtended 30° of visual angle (Fig. 3e, h, k, n). The oculomotor range of marmosets is also more limited than macaques^38^. Thus, the effect of fixation was possibly marginal on the data described above. However, one needs to be cautious when directly comparing retinotopic details of our results with studies that directly controlled fixation.

**Fig. 3.**
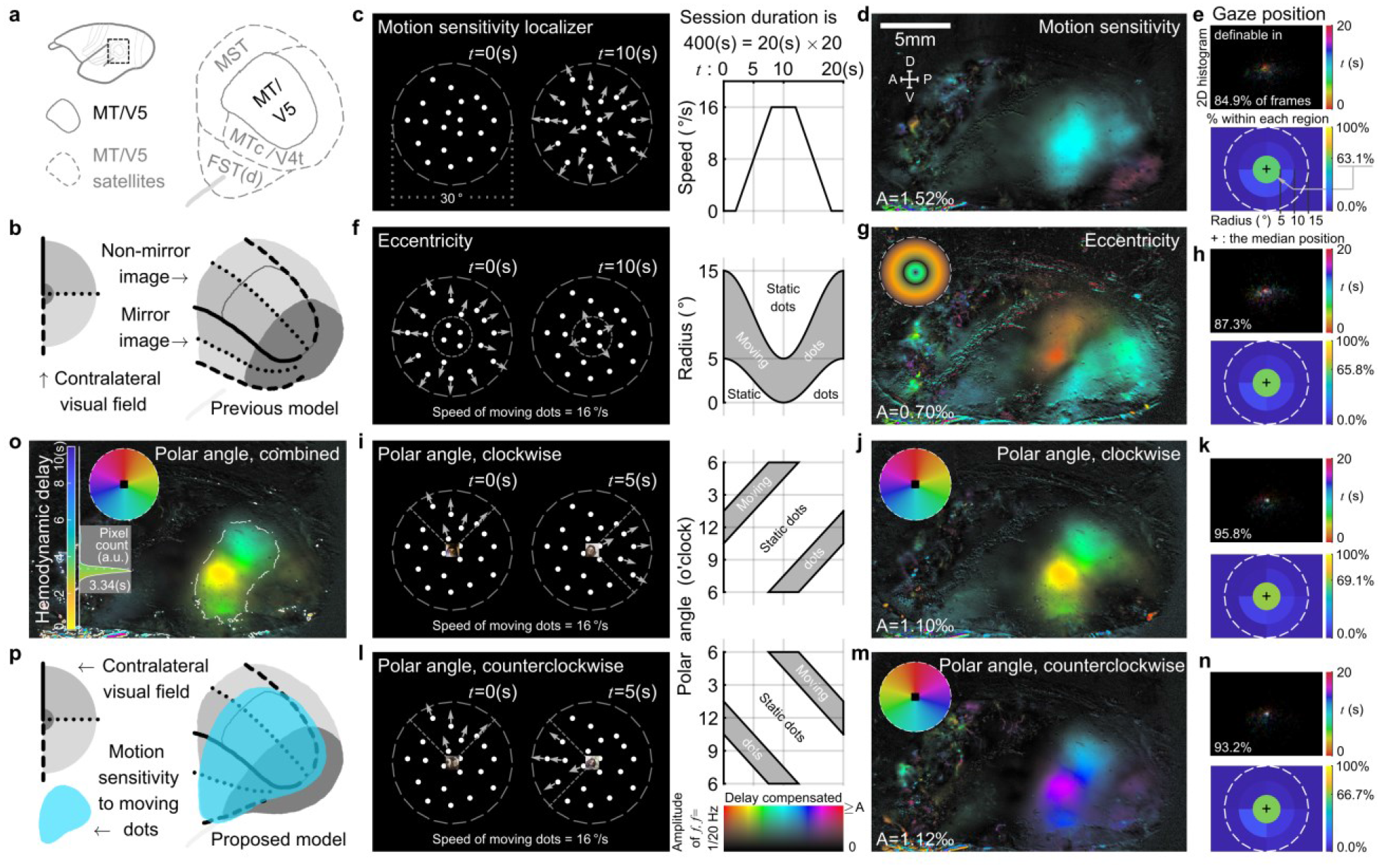
Through-skull retinotopy and motion sensitivity mapping in MT/V5 complex. **a**, A sketch of marmoset MT/V5 complex^9^. MST: medial superior temporal area, MTc/V4t: MT crescent/transitional V4, FST(d): fundal area of the STS (dorsal). **b**, A previous model on the retinotopy of marmoset MT/V5 complex^9, 24^. **c**, Stimuli for motion sensitivity mapping. **d**, Motion sensitivity map. **e**, Eye-tracking results of the motion sensitivity session. **f**, Stimuli for retinotopic eccentricity mapping. **g**, Eccentricity map (wheel: visual field color code). **h**, Eye-tracking results of the eccentricity session. **i**, Clockwise stimuli for retinotopic polar angle mapping. **j**, Clockwise polar angle map (wheel: visual field color code). **k**, Eye-tracking results of the clockwise polar angle session. **l**, Counterclockwise stimuli for retinotopic polar angle mapping. **m**, Counterclockwise polar angle map (wheel: visual field color code). **n**, Eye-tracking results of the counterclockwise polar angle session. **o**, Combined polar angle map (wheel: visual field color code) with a hemodynamic delay histogram. Top 10% of most responsive pixels are outlined by dashed curves and counted in the histogram. Rectangular gray shade: the acceptable delay range. The average delay within this range is 3.34 seconds. **p**, Proposed model for the functional organization of marmoset MT/V5 complex, co-registering the extent of motion sensitivity with the retinotopic map. The motion sensitivity to moving dots extends across two retinotopically organized regions; each has a full representation of the contralateral visual field. Response maps (**d**, **g**, **j**, **m**, and **o**) are shown with tuning phase (color) and amplitude (intensity) separately encoded, according to the 2D colormap in **l** (colormap also transformed into visual field color codes in **g**, **j**, **m**, and **o**). Calibrated gaze positions are definable in 84.9%, 87.3%, 95.8%, and 93.2% of all eye-tracking frames in each session (**e**, **h**, **k**, and **n**). The position of each definable frame is color-coded for its timestamp in the cycle and overlaid in a 2D visual field histogram (top plot). A white spot in the histogram implies the position was not or weakly preferred by any specific time point in the cycle. For each session, the median gaze position, shown as “+” (bottom plot), was estimated 1.1°, 0.9°, 0.8°, and 0.9° away from the very center and 63.1%, 65.8%, 69.1% and 66.7% of gaze positions were within the center 5° radius region (percentage within each designated region is color coded).

### Face patch mapping and a cortical landscape assembled from all through-skull maps

Besides encoding motion, primate visual cortex also processes form and features a specific distributed system for the face (face patches) that has been demonstrated in humans^39–41^, in macaques^36, 39, 42, 43^, and recently in marmosets^44^. These face patches are located along the ventral occipitotemporal cortex and respond to faces much more strongly than to any other object category. Among these face patches, a functional hierarchy is suggested along the posterior-anterior direction with an increase in identity selectivity and view tolerance^42^. A functional specialization is also suggested along the ventral-dorsal direction with an increase in motion preference^40, 43^. One of the face patches described in marmosets, the posterior-dorsal patch (PD), is suggested to locate closely with MT/V5 ^(44)^. This phenomenon has also been reported in macaques^36^ and in humans^41^. Resolving the relative position of this face patch within the local functional landscape including MT/V5 would thus provide new insights into how the face-processing network is organized and has been transformed through evolution. However, the spatial organization of the marmoset PD patch with respect to (i) the local retinotopic map (if any), and to (ii) the extent of motion sensitivity in MT/V5 complex, remains unclear^44^. We sought to fill these gaps by further imaging and contrasting the responses to eight different categories of images (Fig. 4a-c and Extended Data Fig. 8: faces, body parts, animals, fruits & vegetables, familiar objects, unfamiliar objects, spatially scrambled faces, and phase-scrambled faces). A clear patch located at the ventral part of the motion-sensitive region showed a stronger response to faces than to any other control categories (Fig. 4d-g). Additionally, combining all maps provides evidence for a multi-modal functional landscape (Fig. 4h). The scrambling-sensitive region (contrasting faces to scrambled faces, Fig. 4f) mostly coaligned with the center-preferring retinotopic region (Fig. 4h), with a saddle at the location consistent with the foveal region of MT/V5. This saddle is also consistent with relative insensitivity to form in MT/V5 ^(9, 32)^. Furthermore, the face patch (contrasting faces to other objects, Fig. 4g) was located within the motion-sensitive retinotopic region anteroventral to MT/V5, with a center bias and a tendency to prefer the lower visual field. These claims hold for all tested animals (Extended Data Fig. 9). Considering the sensitivity to motion, to face, and to the lower part of the visual field, this face patch may mediate processing of mouth movement. Interestingly, very close to this face patch (Fig. 4h, 2l), a part of the newly discovered tonotopic gradient represents frequencies found in marmoset vocalizations^45^ (∼3-11 kHz). Thus, these vicinal regions may potentially be involved in coordinating audiovisual information in vocal communication.

**Fig. 4.**
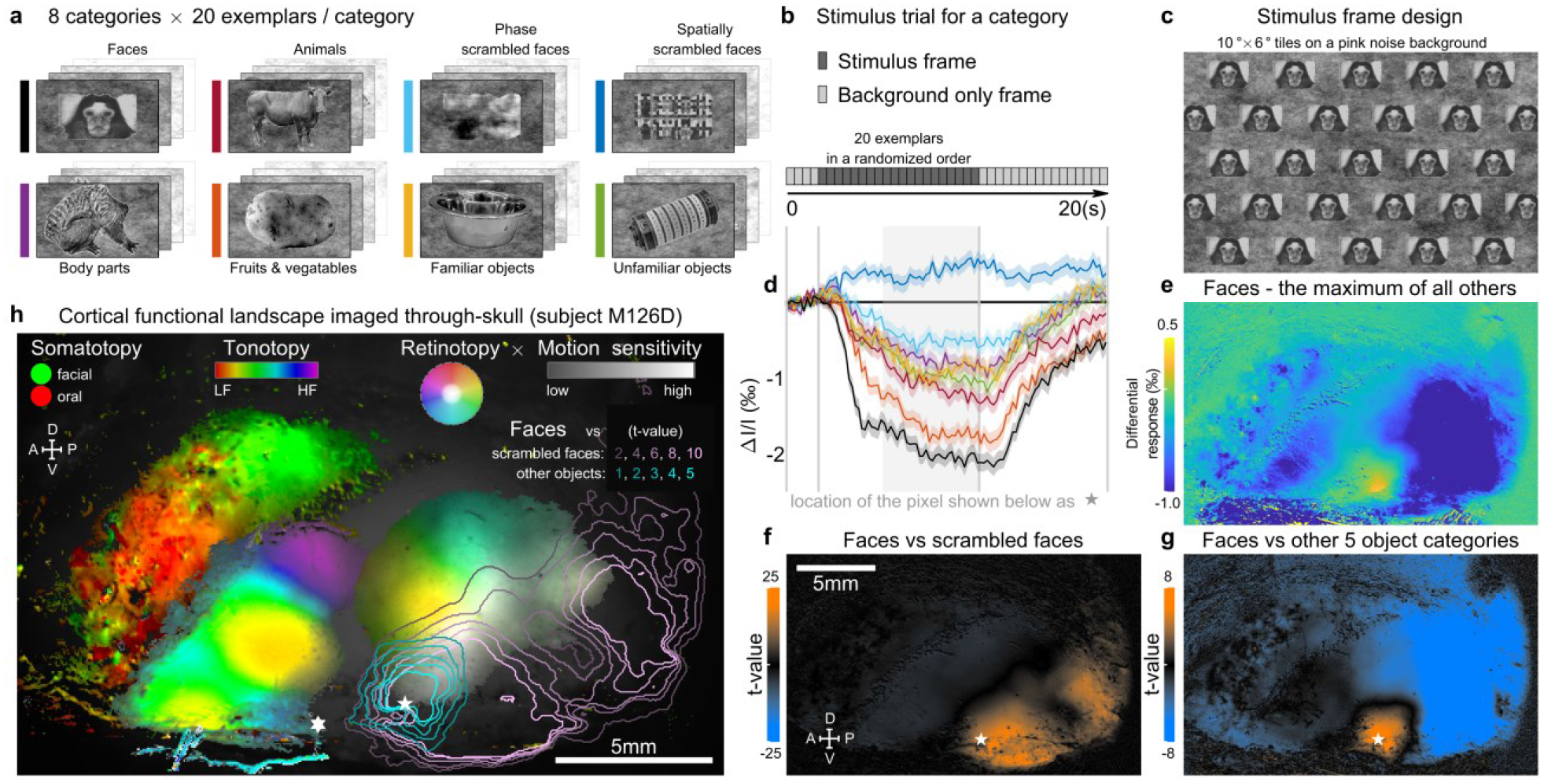
Face patch mapping and a cortical functional landscape assembled from all through-skull maps. **a**, 8 categories of visual stimuli used for face patch mapping (with an exemplar from category shown), including faces^44^, 5 other object categories (body parts, animals, fruits & vegetables, familiar objects, and unfamiliar objects), and 2 types of scrambled faces (spatially, and phase scrambled faces). **b**, The stimulus trial design for measuring the response to a category. **c**, The stimulus frame design for each exemplar (using a face as an example here). The exemplar was tiled in the frame as a stretcher bond pattern. Considering the marmoset oculomotor range^38^, the subject cannot fully avoid the object within its center 5° radius visual field. **d**, Response to each category of a face-selective pixel (location labeled by a pentagram in both **f** and **g**). The colors indicate categories, as shown in **a**. Each average response trace is shown as a solid curve with a shade representing the corresponding SEM (n=218). The rectangular background gray shade indicates an averaging time window for calculating a response value for each trial, with signal polarity compensated. **e**, Differential response map between the mean response to the face category and the maximum of mean responses to all other categories (n=218 for each category). A positive value in the map indicates that pixel has a greater response to faces than any of the other 7 stimulus categories. **f**, The t-value map for comparing faces versus scrambled faces. A t-value is calculated for each pixel in the map by comparing its single-trial responses between those to the faces (n=218) and those to the scrambled faces (n=436). Scrambling sensitivity in the map (orange color: high t-values) implicates involvement in visual form processing^49^. **g**, The t-value map for comparing the face category (n=218) versus all other 5 object categories (n=1090). The face-over-object sensitivity in the map (orange color: high t-values) defines the location of a face patch. **h**, The cortical functional landscape summarized based on modality-specific maps and overlaid on a dimmed image of the recording chamber. The motion-sensitive region is shown with retinotopy together in the “HSV” color space (hue [color]: polar angle; saturation [chroma]: eccentricity; value [brightness]: motion sensitivity, also see Extended Data Fig. 7i). The t-value maps are shown by the contour lines. Pentagram: a face-sensitive pixel (same location as in **f** and **g**). The face patch (outlined by the blue contours) is largely overlapped within both the moving-dots-sensitive region and the retinotopic region representing the lower center of the visual field. Hexagram: a pixel from the newly discovered tonotopic gradient (see Fig. 2m) extending from the rostrotemporal field (RT) of the auditory core to the putative parabelt region, tuned to 5.2 kHz (within the marmoset vocalization range of ∼3-11kHz^45^).

## DISCUSSION

Together, our mapping study through the intact skull without thinning revealed the cortical functional landscape composed of multiple functional gradients and landmarks of various modalities in individual marmosets (Fig. 4h, Extended Data Fig. 9). The ability to map cross-modality landscapes opens new opportunities for examining how the primate cortical system is organized and coordinated across multiple modalities to enable our real-world perceptions and behaviors. This through-skull mapping was enabled by a wide-field, polarization-enhanced optical imaging approach. The approach demonstrated in marmosets here may also find applications in other species, with or without skull thinning. For example, squirrel monkeys, owl monkeys, and some Old-World monkeys (e.g. talapoins^6^) also have skulls that are not much thicker than marmosets (Fig. 1a). The intrinsic optical signal of the brain, however, is inherently limited by hemodynamic change, which may not be fully predicted by local neural signals^46^. Any significant vasculature between the brain and the imaging setup may also affect imaging quality, such as large blood vessels in the dura, diploic veins in the skull, and tissue growth on top of or within the skull (bone degradation). Despite these limitations, the XINTRINSIC system developed in our study could provide data to guide subsequent electrophysiology recordings, two photon imaging, or perturbation experiments that require skull opening. It may yet prove powerful in concert with non-invasive viral gene delivery in primates^47^ for potential through-skull calcium imaging^2, 3^ as well as minimally invasive optogenetic perturbations^48^ where targeting of causal manipulations could benefit from our non-invasive imaging approach. We provide open access to our imaging setup design and software code for straightforward duplication (https://x-song-x.github.io/XINTRINSIC/). With its simple architecture, XINTRINSIC is generally affordable (<$10,000), and has potential for future miniaturization to map cortical functions through-skull in freely roaming primates.

## METHODS

### Through-skull imageability comparison across species

The gyrification index^7, 50^ (GI) and the skull thickness were compared across species based on the data compiled from multiple sources (Fig. 1a). Human (*Homo sapiens*), rhesus macaque (*Macaca mulatta*), common marmoset (*Callithrix jacchus*), loris (*Nycticebus coucang*), lemur (*Cheirogaleus medius*), rabbit (*Oryctolagus cuniculus*), cat (*Felis catus*), and ferret (*Mustela putorius furo*) were reported to have GI values of 2.56, 1.75, 1.18, 1.29, 1.11, 1.15, 1.58, and 1.63, respectively^7^. Cynomolgus monkey (*Macaca fascicularis*), squirrel monkey (*Saimiri sciureus*), owl monkey (*Aotus trivirgatus*), galago (*Otolemur garnetti*), tree shrew (*Tupaia glis*), guinea pig (*Cavia porcellus*), rat (*Rattus norvegicus*), and mouse (*Mus musculus*) were reported to have GI values of 1.65, 1.57, 1.36, 1.25, 1.04, 1.04, 1.04, and 1.02, respectively^50^. All skull thickness data were taken near the location of parietal eminence. Human, rhesus macaque, cynomolgus monkey, common marmoset, squirrel monkey, owl monkey, and loris data were acquired from one primate study as 5.87, 1.52, 1.24, 0.48, 0.62, 0.56, and 1.01 mm, respectively^6^, whereas rabbit^51^, rat^51^, mouse^52^, cat^53^, and ferret^54^ data were 1.50, 0.71, 0.21, 2.90, and 1.20 mm, respectively. Galago (http://dx.doi.org/10.17602/M2/M33809), lemur (ark:/87602/m4/M12869), tree shrew (http://dx.doi.org/10.17602/M2/M6393), and guinea pig (http://dx.doi.org/10.17602/M2/M47035) data were assessed based on the cranium CT scan datasets accessed and archived on MorphoSource.org. The resulting values are 1.48, 0.43, 0.21, and 1.01 mm. The common marmoset used in the present study has a relatively flat brain and thin skull with a GI value of 1.18 (considered lissencephalic^8^) and a skull thickness of 0.48 mm (considered as imageable through-skull). We further confirmed marmoset skull thickness near the location over the auditory cortex by clamping a micrometer onto the edge of acutely prepared bone chips (which may slightly overestimate the thickness due to the skull curvature). The average value is 0.61 mm (n=4), very close to the value reported on the parietal location.

### Marmoset head-cap implant and imaging chamber design

Six marmosets were used in the current study (Extended Data Table 1), including one female, five males, three left hemispheres, and three right hemispheres. The right hemisphere view was mirrored to match the view of the left hemisphere in the figures for display purposes. The subjects’ ages ranged from 44 to 81 months old during imaging acquisition. The basic design of the marmoset chronic head-cap implant has been described previously^55^. Marmosets were adapted to sit calmly in a Plexiglass restraint chair through an acclimation period of two to four weeks. Using sterile techniques, two stainless steel head posts were attached to the skull with dental cement material while the animal was anesthetized with isoflurane (0.5-2.0%, mixed with pure oxygen). During the implant surgery, the lateral part of the skull, presumably over the auditory cortex, the MT complex, and the lateral part of anterior parietal cortex (Fig. 1b) was exposed and covered with a thin layer (∼1 mm) of dental cement. Before applying the dental cement, the putative lateral sulcus was vaguely visible through-skull when the skull was still wet and was thus marked for later reference (water may partially match the refractive index of the skull). The rest of the exposed skull was covered with a thicker layer of dental cement and a wall was formed around the recording chamber to increase the mechanical stability of the head-cap and to protect the chamber. After the animal fully recovered from the implant surgery, the head of the animal was fixed via head posts (Fig. 1c), and the surface of the recording chamber was further polished with acrylic polishing bits (Dedeco, 7750). The dental cement at the chamber bottom optically concealed the surface texture of the skull, presumably by matching its refractive index. We tested two different dental cement materials (orthodontic resin from DENTSPLY Caulk, and C&B Metabond from Parkell [clear]). We did not observe a significant difference between these two materials in terms of imaging performance. Before each imaging experiment, a minimal amount of petroleum jelly was applied evenly at the chamber bottom to further augment the optical smoothness of the surface. The optical axis of the imaging setup was aligned as perpendicular as possible to the local surface. The animal was head-fixed and kept awake through all imaging sessions. Outside imaging, the chamber was protected and sealed with a dental impression material (GC America, EXAMIX NDS injection). All experimental procedures were approved by the Johns Hopkins University Animal Use and Care Committee.

### Imaging setups

The general optical design of XINTRINSIC is illustrated in Extended Data Fig. 1C. A swappable LED (Thorlabs M470L3, M530L4, M590L4, M625L3, M730L5, and M850L3; corresponding to nominal wavelengths [colors] of 470 nm [blue], 530 nm [green], 590 nm [amber], 625 nm [red], 730 nm [far-red], and 850 nm [near inferred, or NIR], respectively) passed light through a Koehler illuminator consisting of 4 lenses (Thorlabs ACL12708U-A/B, Edmund Optics 49-793-INK, 33-923-INK, and 49-372-INK), a field stop, and an aperture stop. The light from the Koehler illuminator was filtered by a polarizer (Thorlabs WP25L-VIS) before being reflected by a polarizing beam splitter ([PBS] Thorlabs, WPBS254-VIS) to a 1× telecentric objective (Thorlabs TL1X-SAP, without the accessory wave plate). The polarizer was aligned to maximize S-polarized light throughput to the PBS. The light power under the objective was measured as not more than 50mW before *in vivo* imaging. Backscattered light was collected by the same objective, transmitted through and filtered by the same PBS, and then passed to another polarizer placed to allow P-polarized (cross-linearly polarized) light to go to the coupling relay lenses (Thorlabs, A508-080-AB-ML, ×2). The light was eventually focused by a camera lens (Navitar, NMV-6X16, at the maximal zoom) onto a high-saturation-capacity, high-frame-rate, mass-produced CMOS camera (FLIR, grasshopper 3, GS3-U3-23S6M-C, with a SONY IMX174 sensor). The imaging acquisition of the camera was controlled by an 80 Hz trigger pulse train generated on a data acquisition (DAQ) card (NI, PCIe-6323). Recordings were acquired with custom code in Matlab (Mathworks, R2018a and later), at a resolution of 1920×1200 pixels and a speed of 80 frames/second (1920×1200@80fps). The data were binned and saved in a format of 480×300@5fps. Based on a Poisson process assumption for incoherent photon arrivals, the theoretical shot noise floor under the current condition over 20 repetitions was estimated as 0.08‰. Besides the Koehler illuminator built in the XINTRINSIC setup, another independent illuminator, coupled with fiber bundles as output ports but without polarization control, was built for control purposes (Fig. 1d). The light from one of the swappable LEDs was passed through four coupling lenses with a diffuser in the middle (Thorlabs ACL12708U-A/B, AC254-045-A-ML, ED1-S20, ACL5040U-A, and ACL50832U-A) to fulfill both the input aperture (4.52mm) and the numerical aperture (NA=0.57) of a gooseneck Y-bundle light guide (Thorlabs, OSL2YFB). The output ports were pointed from the side of the objective to illuminate the chamber as homogenously (to the best of ability).

### Stimulus delivery

For somatosensory stimulation, air puffs were powered by a pneumatic air supply to a set of solenoid valves (SMC, S070M-6DC-40) with a 0.38 MPa pressure at the valve inlet. The outlets of the valves were further routed through 2-mm-inner-diameter 1.5 meters long tubing and 10-30 cm long adjustable hoses (Loc-Line, 1/4”) before spraying air out of nozzles (Loc-Line, 1/16” round nozzle). The airflow intensity out of the nozzle was tested on the experimenter’s skin and confirmed to be appropriately mild before pointing onto the subject. An extra nozzle was placed about 3 cm away from the animal’s ear (contralateral to the imaging chamber), with the airflow directed away from the subject. Air buffs at the same repetition rate were delivered through this nozzle whenever there were no somatosensory puffs delivered from the other nozzles to counterbalance the acoustic noise generated from the other somatosensory stimulating nozzles. All solenoid valves were switched by control signals generated on the same DAQ card that controlled the imaging acquisition.

For auditory stimulation, digital waveforms were converted into analog signals on the same DAQ card (16-bit resolution, 100,000 samples/second conversion rate). The analog signal was attenuated by a programmable attenuator (Tucker-Davis Technology, PA5) before feeding into an amplifier (Crown, D75) that powered a loudspeaker (KEF, LS50), placed ∼1 meter away in front of the subject. Sound levels were calibrated by placing a microphone (Brüel & Kjær, type 4191) at the animal’s typical head position.

For visual stimulation, the display content was controlled by custom code in Matlab, using the Psychophysics Toolbox extension^56^. An LCD monitor (LG, 32GK850F, 32-inch nominal size, 144 Hz refreshing rate, 2560×1440 resolution, 400 cd/m^2^ maximal brightness) was placed ∼75 cm away in front of the subject. An eye-tracking camera (FLIR, FMVU-03MTM-CS; Edmund Optics, 86-410), and a collimated NIR LED (Thorlabs, M850L3 and ACL5040U-DG15-B) were placed right above the monitor to track the animal’s eyes during visual experiments. The eye-tracking camera was controlled by a trigger sequence at 10 fps generated also on the DAQ card.

The air nozzles, the loudspeaker, and the monitor, together with the XINTRINSIC imaging setup and the subject were positioned inside a double-walled soundproof chamber (Industrial Acoustics, custom model), whose interior was covered by 3-inch acoustic absorption foam (Pinta Acoustic, Sonex). Stimulations of different modalities were all synchronized with the imaging acquisition. The somatosensory and auditory stimulations shared the same hardware clock and start trigger on the DAQ card, whereas the DAQ card reset the trial time clock used in Matlab for visual stimuli generation at the beginning of every trial or cycle.

### Experimental design

For the parcellation experiment, the stimulating air nozzle was placed facing the subject and ∼5-8 mm away from the subject’s mouth. The airflow direction was slightly tilted upward to target the regions of the face and the mouth below the nasal and ocular areas. The LCD monitor for visual stimulation was placed between the loudspeaker and the subject. The sound level was re-calibrated to compensate for the effect of the monitor placed in between. A 396-second-long recording session simultaneously consisted of 18 cycles of a 22-second-long somatosensory stimulation trial, 20 cycles of a 19.8-second-long auditory stimulation trial, and 22 cycles of an 18-second-long visual stimulation trial (Fig. 1f). The somatosensory stimuli were a 6-second-long, 10-Hz-repetition-rate air-puff train, presented at the center of each somatosensory trial cycle (Fig. 1f left). To generate each air puff, the solenoid valve was switched on for 0.05 seconds and then off (50% duty cycle). The auditory stimuli were a 9.8-second-long, 5-Hz-repetition-rate white noise burst sequence, presented at the center of each auditory trial cycle (Fig. 1f middle). Each white noise burst was 0.1 seconds in duration, with 20-millisecond sine ramps at the onset and the offset. The burst was 80 dB SPL loud when it was on. The visual stimuli were radially outward moving dots varying in speed (Fig. 1f right). The display background was black, and the dots were white. Each dot was 0.4° in viewing diameter. Initially, 180 dots were randomized in position within a 15°-radius display field. Whenever a dot moved out of this field range, another replacing dot was generated in the very center of the field with a randomized moving direction. During each 18-second-long visual trial cycle, the instantaneous speed of all dots was 0°/second for the first 2 seconds, then linearly increased to 16°/second during the next 5 seconds, stayed at 16°/second for the next 4 seconds, linearly decreased to 0°/second during the next 5 seconds, and stayed at 0°/second for the last 2 seconds of the cycle. Simultaneous eye-tracking was performed to make sure the subject was alert and generally kept the gaze in the center.

For the somatotopy mapping experiment, two stimulating nozzles were configured. The oral nozzle was placed to approximate the subject’s incisors, whereas the facial nozzle was placed to target the cheek contralateral to the imaging side but kept ∼5 to 8 mm away from the subject (Fig. 2b). A 437-second-long recording session simultaneously consisted of 19 cycles of a 23-second-long oral stimulation trial and 23 cycles of a 19-second-long facial stimulation trial (Fig. 2c). The oral stimuli were a 4-second-long, 10-Hz-repetition-rate air-puff train presented at the center of each oral trial cycle (Fig. 2c left), whereas the facial stimuli were another 4-second-long 10-Hz-repetition-rate air-puff train but presented at the center of each facial trial cycle (Fig. 2c right). Two parallel solenoid valves were routed together to power the facial nozzle. To generate each air puff, the corresponding solenoid valves were switched on for 0.03 seconds and then off (30% duty cycle).

For the tonotopy mapping experiment, two sessions were recorded to derive the tonotopic map in each subject. Each session was 400 seconds long and consisted of 20 cycles of a repeating 20-second-long trial (Fig. 2i). The trial started with 2.7 seconds of silence, followed by a sequence of 73 pure tone pips, and ended with another 2.7 seconds of silence. Each pure tone pip was 0.2 seconds in duration and had 20-millisecond sine ramps at both the onset and the offset. All pure tone pips were played at 50 dB SPL loudness. The “downward” session (Fig. 2i left) had the trial cycle with the tone pip sequence decreasing in frequency for 6 octaves (72 semitones), starting with a pure tone pip of 28,160 Hz (music note A10), continuing with each of the following pips descending one semitone in frequency from the previous pip, and ending with a pure tone pip of 440 Hz (A4). The “upward” session (Fig. 2i right) was the opposite and had the trial cycle with the pure tone pip sequence starting with a pure tone pip of 440 Hz (A4) and ending with a pure tone pip of 28,160 Hz (A10).

For the motion sensitivity and retinotopy mapping experiment, four sessions were recorded in each subject. All sessions were designed based on 0.4°-diameter white dots moving radially outward within a 15°-radius display field and on a black background^57^. Each session was 400 seconds long and consisted of 20 cycles, each 20 seconds long. The first session was to re-map the spatial extent of motion sensitivity to moving dots (Fig. 3c), and the design was very similar to the visual part of the parcellation experiment. During each cycle, the instantaneous speed of all dots was 0°/second for the first 2 seconds, then linearly increased to 16°/second during the next 6 seconds, stayed at 16°/second for the next 4 seconds, linearly decreased to 0°/second during the next 6 seconds, and stayed at 0°/second for the last 2 seconds of the cycle (Fig. 3c right). The 16°/second speed of moving dots is very close to the median of preferred speeds to moving dots by marmoset MT neurons^58^. The second session was to map the retinotopic eccentricity (Fig. 3f). The dots moved outward at a speed of 16°/second, within a range determined by eccentricity, and otherwise stayed static. The inner and the outer limits of the allowed motion range were both sinusoidally modulated and synchronized with the 20-second-long cycles at a cosine phase. The outer limit started at a 15°-radius at the beginning of each cycle and dropped to 5° at the middle of each cycle, whereas the inner limit started at a 5°-radius at the beginning of each cycle and dropped to 0° at the middle of each cycle (Fig. 3f).

Thus, at the beginning of each cycle, only the periphery of the field (5°-15° radius) contained moving dots, leaving the center of the field (0°-5° radius) with static dots (Fig. 3f left), whereas at the middle of each cycle, only the center contained moving dots, leaving the periphery with static dots (Fig. 3f right). The third (Fig. 3i) and the fourth (Fig. 3l) sessions were to map the retinotopic polar angle tuning. The dots moved outward at a speed of 16°/second, only within a range determined by polar angle, and otherwise stayed static. This motion range was a quarter circle in shape, with its middle line pointing at the 12 o’clock polar angle at the beginning of each cycle. The shape of this range persisted while the angle at which the range middle line pointed was swept either clockwise (Fig. 3i) or counterclockwise (Fig. 3l) for each session. The sweeping speed was 0.05 rounds/second and was thus synchronized with the 20-second-long cycles. Different from the first two sessions, in which the motion range patterns at any moment were always rotationally symmetric and thus did not carry a natural bias to drive the subject’s attention to any specific polar angle at any moment, the motion range pattern of these polar angle testing stimuli was not rotationally symmetric at a given moment and may carry such an attention bias. To draw the subject’s attention to the center, we displayed random small marmoset pictures (∼3°×3°) in the center of the field on the top of the dots during these polar angle testing sessions. Each marmoset picture lasted for 2 seconds. These pictures were acquired through Google Images and further cut and scaled to the desired size. Simultaneous eye-tracking was performed during all sessions (see below for details). Indeed, the eye-tracking results showed the subject’s gaze was kept predominantly to the center of the field in these polar angle sessions (Fig. 3k and n), with 69.1% and 66.7% of total detected gaze positions falling in the center 5°-radius range, compared to 63.1% and 65.8% in the first two sessions (Fig. 3e and h).

For the face-patch mapping experiment, six categories of visual objects were tested: marmoset faces, marmoset body parts, animals, fruits and vegetables, familiar objects, and unfamiliar objects. The images of marmoset faces were obtained from the authors of the pioneer marmoset face-patch mapping study^44^. The images of familiar objects were taken in our colony and laboratory at Johns Hopkins. The images of other categories were searched and selected using Google Images. Visual objects were extracted from the original images, converted to grayscale, and further equalized for contrast and average luminance^59^. The objects were scaled and rotated to minimize the variation in aspect ratio and to minimize the variation across categories for the mean value and the standard deviation of object size (defined as the area each object covers) within each category. Two additional categories of scrambled faces were generated. For both spatially and phase scrambled faces, the same contour mask of the corresponding face was applied back to the scrambled faces so that the original contour edges remained. Together, 20 exemplar objects were generated for each of the 8 categories (Fig. 4a and Extended Data Fig. 8). To synthesize a stimulus frame for each object, the object was repeated and tiled as a stretcher bond pattern in the frame (Fig. 4c). Each “tile” was about 10°×6° in size (480×288 in display pixel). Considering the range of marmoset gaze is limited and is no larger than the stimulus frame displayed^38^, no matter where the subject gazed at, it could not fully avoid the object within the 5°-radius center of its visual field. The background of the frame was set as a randomly generated pink noise image that matches the average luminance of the objects (115 on 8-bit grayscale). To test the cortical response to a category, a 20-second-long trial was generated online with 40 display frames (Fig. 4b). Each frame lasted for 0.5 seconds. The first 4 frames were with pink noise background only, followed by stimulus frames of all 20 exemplar objects of the category in a randomized order, and then ended with another 16 pink noise background only frames. Stimulus trials of these 8 categories were presented in a randomized order for each testing cycle. A testing block contained 2 testing cycles and was therefore 320 seconds long. A random music selection from spotify.com was played through the loudspeaker during testing blocks to keep the subject alert. The subjects rested between testing blocks. On the first day of the experiment, a reference image of the imaging field of view was taken. On each following day, this reference image was displayed simultaneously with the real-time image (through different color channels) to align the images and maintain a consistent field of view. Testing blocks acquired on different days were then pooled together for analysis. Simultaneous eye-tracking was performed with the imaging acquisition.

### Imaging analysis

A phase-encoding strategy^18, 22,30^ was utilized in designing all these experiments except for the face-patch experiment. To analyze phase-encoding experiments, the trace of each pixel across the entire recording session was Fourier transformed and then normalized to intensity change over baseline intensity (ΔI/I). Signal polarity, defined as whether a positive or negative change in ΔI/I indicates a higher response to a stimulus, varies across wavelengths and was empirically determined for each wavelength from the activation patterns of all three tested modalities (Extended Data Fig. 2) as negative for 470 nm (blue), negative for 530 nm (green), ambiguous, presumably positive for 590 nm (amber), positive for 625 nm (red), positive for 730 nm (far-red), and negative for 850 nm (NIR). This difference in polarity can be explained by the difference in how much each wavelength emphasizes changes in oxy-hemoglobin (HbO) over changes in deoxy-hemoglobin (HbR) during a hemodynamic response (see Extended Data Fig. 2 and Extended Data Fig. 1b). Signal polarity was compensated to calculate each pixel’s phase response to the stimulus at the corresponding frequency.

Research has suggested that hemodynamic response begins within ∼0.5 seconds of stimulus onset and peaks at 3-5 seconds after the onset^60^. To decode the preferred stimulus phase (or tuning phase) from the raw phase in response, it is necessary to compensate for this hemodynamic delay. For both the tonotopy mapping and the retinotopy polar angle mapping experiments, two sessions were performed with stimuli that were reversely identical to each other. The “upward” pure tone pip sequence was the time-reversal of the “downward” pure tone pip sequence, whereas the clockwise polar angle sweep was the time-reversal of the counterclockwise polar angle sweep. Thus, by assuming the hemodynamic delays of a pixel in these two temporally “mirrored” sessions are identical^18^, this delay can be numerically derived or canceled out by reversing the response phase of one session and combining it with the response phase of the other session (Fig. 2i and j). We considered the top 10% of pixels, ranked by their mean response amplitudes at the corresponding frequency in these two sessions, as candidate pixels to estimate the hemodynamic delay. These pixels were color-coded for the derived hemodynamic delay in each subject’s delay map (Fig. 2k, 3o, Extended Data Fig. 5g, 7d). The delay values of these pixels were counted and shown in a histogram with the same color-code (Fig. 2k, 3o, Extended Data Fig. 5h, 7e). These delay values largely fell within a range of 2.0-5.7 seconds. A peak of this range in the histogram was located within a narrowed range of 3.2-4.4 seconds. By assuming 2.0 to 5.7 seconds as the empirically acceptable delay range, we further derived the average hemodynamic delay in each subject among qualified pixels. For the tonotopy experiment, five tested subjects each had an average delay of 3.94, 3.61, 3.47, 4.46, and 4.12 seconds, with a group average of 3.9 seconds (Extended Data Fig. 5h). For the retinotopy polar angle experiment, five tested subjects each had an average delay of 3.34, 3.42, 3.72, 2.96, and 3.91 seconds, with a group average of 3.5 seconds (Extended Data Fig. 7e). Thus, to decode the tuning phase from the raw response phase for a single session, we compensated 3.9 seconds for the hemodynamic delay in tonotopy tuning phase analysis (Extended Data Fig. 5), 3.5 seconds in retinotopy polar angle tuning phase analysis (Extended Data Fig. 7), and 3.7 seconds in all other single-session tuning phase analyses.

The tuning maps were drawn after compensating for signal polarity and delay at the corresponding frequency for each pixel. The HSV (hue-saturation-value) color space was utilized to visualize both responsiveness and tuning for each pixel. The tuning phase was encoded in the hue (color) channel, whereas the response amplitude of the corresponding frequency was encoded in the saturation (chroma) and value (brightness) channels. Since the saturation and value channels can only have values between 0 and 1, for most of the maps, we encoded the response amplitude at the 99th percentile of all pixels as the upper limit value 1 in both saturation and value channels (Fig. 1h, 2d, 3d, 3g, 3j, 3m, 3o, Extended Data Fig. 2, 3, 4, 7), except for the amber light in Extended Data Fig. 2 (97th percentile), the subject M117B in Extended Data Fig. 3 (97th percentile), and Extended Data Fig. 5 (95th percentile). The upper display limit of response amplitude was set at absolute values as indicated in Fig. 2k, 2l, and Extended Data Fig. 6. For tonotopy visualization, pixels that had their tuning phases outside the time range of the tone pip sequence were set to 0 in their saturation channel. To visualize the retinotopy tuning together with the motion sensitivity, we encoded retinotopy polar angle tuning in the hue (color) channel, retinotopy eccentricity tuning in the saturation (chroma) channel, and motion sensitivity in the value (brightness) channel (Fig. 4h, Extended Data Fig. 7i). To generate a summary map for a session designed with multiple stimulation frequencies (e.g. parcellation, somatotopy experiments), the RGB (red-green-blue) color space was utilized to display maps of different frequencies together (Fig. 1i, 2e, Extended Data Fig. 2e, 3e, 4d). For the map of each stimulation frequency, a mask was first generated with pixels that had tuning phases falling into the trial time range corresponding to the most intense stimuli with 1-second tolerance at either side. Pixels included in this mask had their amplitudes in the original HSV “value” channel further shown in the designated RGB color channel (red, green, or blue).

For the face-patch experiment, 218, 134, and 234 testing cycles were recorded in the subjects M126D, M15E, and M117B, respectively. The pixel response trace of each trial was normalized to the pre-stimulus baseline, defined as the average signal value of the first 2 seconds in the trial (Fig. 4d, Extended Data Fig. 9b, 9g, 9l). A single ΔI/I response value was extracted for each trial by averaging the normalized and polarity-compensated response within a time window of 4 to 10 seconds after the first stimulus frame onset. Two t-value maps were drawn for each subject. One map was to compare response values between those to the face category and those to all other five object categories combined (Fig. 4g, Extended Data Fig. 9e, 9j, 9o). The other map was to compare response values between those to the face category and those to the two scrambled face categories combined (Fig. 4f, Extended Data Fig. 9d, 9i, 9n).

To put maps of different experiments together (Fig. 4h, Extended Data Fig. 9f, 9k, 9p), somatotopy, tonotopy, and retinotopy of the same subject were co-registered to and overlaid on top of a chamber bottom image (also dimmed and served as the display background). Maps of t-values were also co-registered to the same map but shown as iso-t-value contour lines. Together these maps formed a cortical functional landscape in each subject.

### Eye-tracking

To determine the absolute position of the subject’s gaze, calibration sessions were performed in addition to other visual experiments performed during each day of visual experiments. 80 marmoset images were acquired from Google Images. Each image was cut and scaled to a size of 160×160 pixels. The screen was divided into a 15×9 location grid to display these images without overlap and with one right in the center. Among these 15×9 locations, 37 locations had their centers within the center 23° diameter range of visual angle. An image was displayed at one of these 37 locations for 1.5 seconds, with the rest of the monitor displaying a gray background. 80 images were displayed in a pseudorandomized order. Eye-tracking recording from this session (120 seconds in duration) was later used for calibrating the absolute gaze positions in the other experimental sessions. This calibration grid generally covers the entire marmoset oculomotor range, which is more limited than that of macaques^38^.

## DATA AVAILABILITY

The data that support the findings of this study are available from the corresponding author upon reasonable request.

## CODE AVAILABILITY

Data collection routine was performed with custom MATLAB code, available at https://github.com/x-song-x/XINTRINSIC (the commit on Dec, 9, 2019), and https://github.com/x-song-x/ChKshared (the commit on Dec, 7, 2019). Data analysis routine was performed with custom MATLAB code, available at https://github.com/x-song-x/fluffy-goggles (the commit on Aug 25, 2020).

## ACKNOWLEDGEMENTS

This research was supported by National Institute of Health grants DC003180 and DC005808 to X.W. X.S. was supported by a fellowship from the Kavli Neuroscience Discovery Insitute at JHU. We thank S. H. Park and D. Leopold for providing marmoset face images, J. Lynch, K. Schonvisky, S. Miller, and J. Izzi for assistance with animal care, Q. Fang for technical advices on the MCXLAB simulation toolbox, and E. Issa, X. Liu, L. Zhao and M. Osmanski for their comments on the earlier versions of the manuscript. Creation of datasets accessed on MorphoSource was made possible by the following funders and grant numbers: NERC NE/G001952/1 to N.S. Jeffery, NSF BCS 1304045 and a research grant from Trinity College of Arts and Sciences to C. Wall, and NSF BCS 1540421 to G.S. Yapuncich and D.M. Boyer.

## AUTHOR CONTRIBUTIONS

X.S. and X.W. designed the study. X.S., Y.G., and X.W. developed the marmoset preparation. X.S. developed the XINTRINSIC setup. Y.G. performed the Monte Carlo simulation. C.C. and X.S. designed the pilot through-skull testing. H.L. and X.S. designed the eye-tracking system and the stimuli in the face-patch mapping experiment. X.S. designed the mapping experiments and analyzed the data. X.S., Z.S., and Y.G. performed the mapping experiments. X.S., Y.G., and X.W. wrote the manuscript with input from all authors. X.W. supervised the study.

