## Supplementary Figures for "Functional maps of the primate cortex revealed by through-skull wide-field optical imaging"

### EXTENDED DATA FIGURES

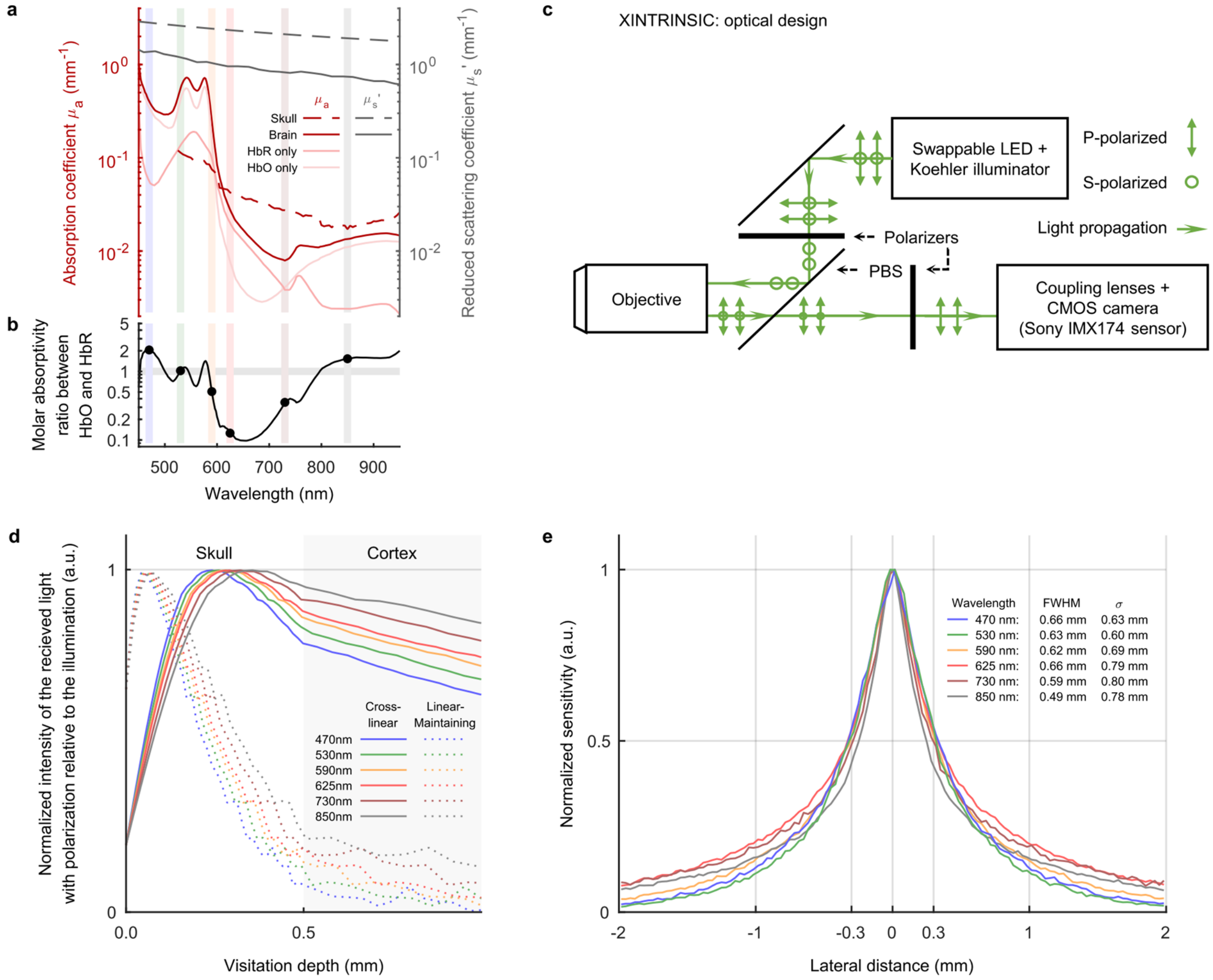


**Extended Data Figure 1.**

**Optical analysis of through-skull intrinsic signal and the imaging strategy. a**, Absorption and scattering properties of the skull and the brain. The red curves indicate absorption coefficients ($\mu_a$), whereas the gray curves indicate reduced scattering coefficients ($\mu'_s$). The coefficients of the skull are shown in dashed curves, whereas the coefficients of the brain are shown in solid curves (see Supplementary Information for data sources and calculation). Vertical color stripes indicate LED wavelengths tested in the current study and their corresponding colors. In both skull and brain, scattering is the dominant effect (coefficients are higher) over absorption. The absorption of the brain is much stronger at shorter wavelengths (blue and green) than at longer wavelengths (red, far-red, and NIR). Absorptions of the HbR and HbO components in the brain are separately drawn as pale red curves (assumed composition of the blood: 25% HbR, 75% HbO). **b**, The molar absorptivity ratio between HbO and HbR (see Supplementary Information for data sources and calculation). This ratio varies across different wavelengths (shown by the black solid curve). The ratios at the wavelengths tested in the current study are marked by filled black circles. The gray horizontal strip represents an isosbestic line. The intersection points between this line and the ratio curve indicate that the corresponding wavelengths are absorbed equally by HbR and HbO at the same molar concentration. The green light (530 nm) used in the current study has a molar absorptivity ratio close to the isosbestic line, suggesting the intrinsic signal recorded with the green light measures changes in total hemoglobin concentration independently from changes in blood oxygenation[3]. In contrast, the blue and the NIR lights emphasize HbO, whereas the amber, the far-red, and the red lights emphasize HbR. **c**, A sketch of the optical design of XINTRINSIC. Light polarization directions and light propagation directions are labeled. **d**, An estimation of photon visitation depths, modified from a previous Monte Carlo simulation[21] (see Supplementary Information), in which a linearly polarized beam was shed perpendicularly into the “tissue”. Visitation depth was defined as the maximum depth that a photon visits in the tissue before being backscattered out of the “tissue”. The photons that were depolarized and then collected from the cross-linear polarization channel visited deeper structures than the photons that maintained their initial linear polarization. These “cross-linear” photons thus are presumably more sensitive to hemodynamic changes in the cortex. The intensity was individually normalized by the peak of each curve. **e**, A lateral resolution estimation of through-skull imaging signal for picking up cortical hemodynamic changes in marmosets, from our own Monte Carlo simulation (see Supplementary Information). The intensity was individually normalized by the peak of each curve. Estimated FWHM and standard deviation ($\sigma$) of the lateral profile at each wavelength are, respectively, 0.66 and 0.63 mm for 470 nm (blue), 0.63 and 0.60 mm for 530 nm (green), 0.62 and 0.69 mm for 590 nm (amber), 0.66 and 0.79 mm for 625 nm (red), 0.59 and 0.80 mm for 730 nm (far-red), and 0.49 and 0.78 mm for 850 nm (NIR).

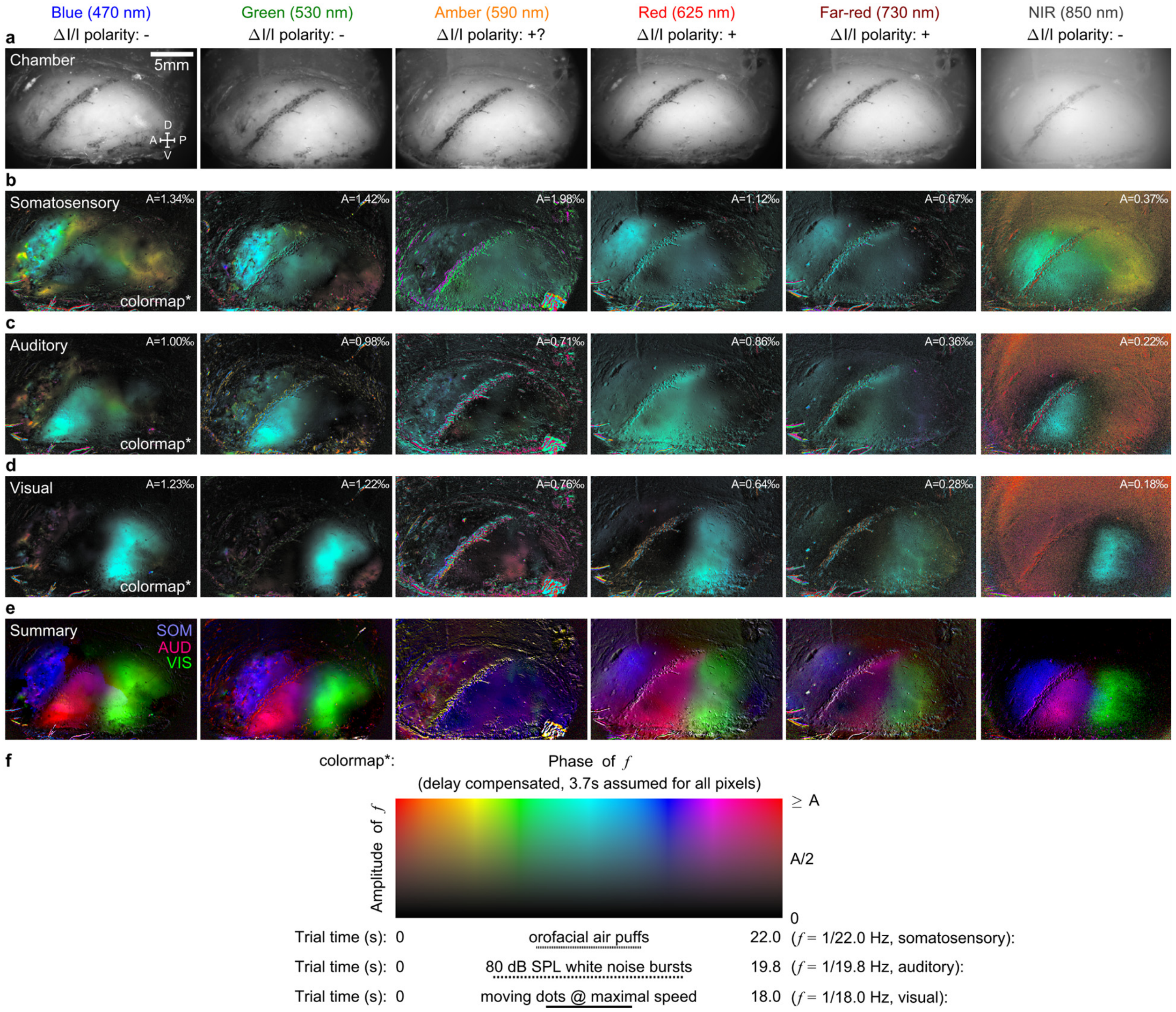


**Extended Data Figure 2.**

**The effect of wavelength on modality parcellation.** The modality parcellation experiment presented in Fig. 1 was performed in subject M126D with six different illumination wavelengths, which are shown here in six columns. The color, the wavelength, and the assumed signal polarity (the direction of light intensity change in response to stimulus) of each illumination light are indicated on the top of each column. **a**, The image of the recording chamber taken at each wavelength. **b**, The somatosensory response map under each wavelength. **c**, The auditory response map under each wavelength. **d**, The visual response map under each wavelength. All response maps (**b-d**) are compensated for signal polarity according to the assumed signs in the titles. A fixed hemodynamic delay value of 3.7 seconds is assumed and compensated for all pixels in these maps. Response phase and amplitude are visualized separately as color and intensity in these maps, according to the 2D colormap in **f**. For each response map, the upper display limit of amplitude "A" is listed at the top right. **e,** A summary map for each wavelength, with SOM on the blue color channel, AUD on the red color channel, and VIS on the green color channel. **f,** The 2D colormap for the response maps shown in **b**, **c**, and **d**.

For all three modalities, the shorter wavelengths (blue and green) had superior response amplitudes over other wavelengths. This is consistent with the amplitude of the absorption spectra - the blood in the brain absorbs the shorter wavelengths (blue and green) much more efficiently than other wavelengths (Extended Data Fig. 1a). Although a longer wavelength (e.g. NIR) would have weaker scattering and absorption and thus penetrates deeper brain structures, the signal amplitudes were much smaller.

The response patterns of each modality were generally consistent among most of the wavelengths and agreed with the atlas (Fig. 1b), confirming the assumed signal polarities are valid for most of the wavelengths. The observed signal polarity difference among wavelengths is correlated with and can be explained by the HbO/HbR molar absorptivity ratio (Extended Data Fig. 1b). Any wavelength with the ratio higher than 0.5 (blue, green, and NIR) showed a negative signal polarity, whereas any wavelength with the ratio lower than 0.5 (red and far-red) showed a positive signal polarity. The amber color with a ratio of ~0.5 showed ambiguity in signal polarity. It was suggested the major hemodynamic response following a stimulus consists of an HbO increase and an HbR decrease[60]. The wavelengths with higher ratios (blue, green, and

NIR) would pick up the HbO increase more over the HbR decrease and are thus more absorbed during the hemodynamic response (negative ΔI/I), whereas the wavelengths with lower ratios (red and far-red) would pick up the HbR decrease more over the HbO increase and are thus less absorbed during the hemodynamic response (positive ΔI/I). The amber light may pick up these opposite changes more equally and is thus ambiguous in signal polarity.

Different from the somatosensory and the visual response patterns, which are consistent across all measurement wavelengths (except for the amber light), the auditory response patterns measured with the higher-ratio wavelengths (green, blue, and NIR) generally agree with each other but are different from the patterns measured with the lower-ratio wavelengths (red and far-red). Many previous attempts using intrinsic signal to map the auditory cortex in a variety of species (chinchillas[61], cats[62, 63], ferrets[64, 65]) only succeeded with green wavelengths but failed with red wavelengths. These studies, together with our data here, imply that the HbR hemodynamic response may behave differently in the auditory cortex than in the other cortical areas in a variety of species[61-65]. Further investigations are needed to reveal more detailed hemodynamic similarities and dissimilarities in different cortical areas and species.

Considering the signal amplitude, the ambiguity in signal polarity, and the consistency in mapping auditory cortex, the shorter wavelengths (blue and green) are more suitable for mapping cortical hemodynamic responses through the intact skull in awake marmosets.

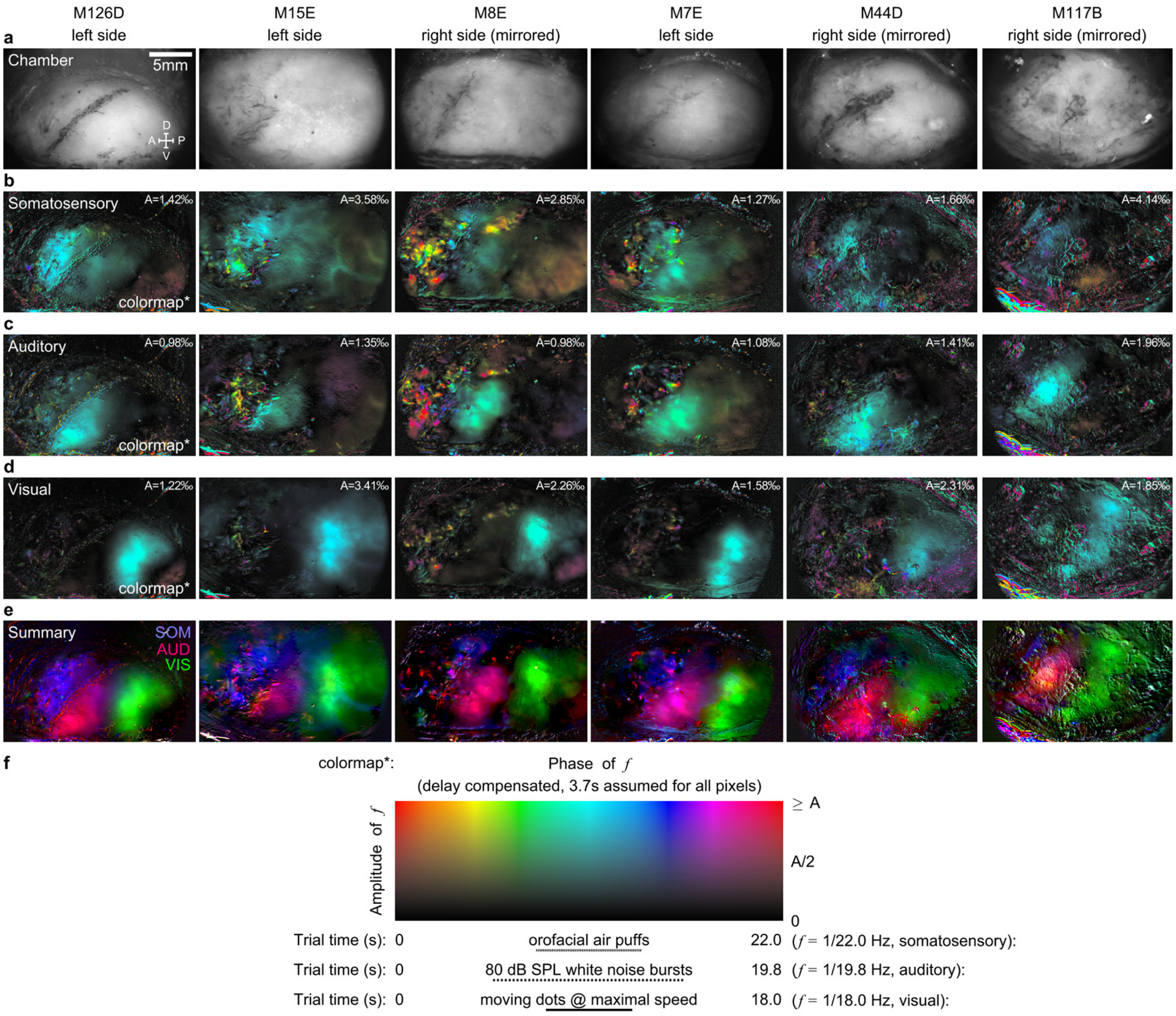


**Extended Data Figure 3.**

**Modality parcellation in six tested subjects.** The modality parcellation experiment presented in Fig. 1 was performed in six subjects which are shown here in six columns. Subject ID and the imaged hemisphere are indicated on the top of each column. The right hemisphere view is mirrored to match the view of the left hemisphere for display purposes. **a**, The image of the recording chamber in each subject. **b**, The somatosensory response map in each subject. **c**, The auditory response map in each subject. **d**, The visual response map in each subject. A fixed hemodynamic delay value of 3.7 seconds is assumed and compensated for all pixels in these maps (**b-d**). Response phase and amplitude are visualized separately as color and intensity in these maps, according to the 2D colormap in **f**. For each response map, the upper display limit of amplitude "A" is listed at the top right. **e**, A summary map for each subject, with SOM on the blue color channel, AUD on the red color channel, and VIS on the green color channel. These response patterns are generally consistent with the atlas (Fig. 1b). Subject M117B had a chamber that barely covered any part of the somatosensory cortex and had part of the skull (~3 mm × 4 mm) over the auditory cortex thinned in an earlier pilot experiment. Comparing to the other subjects, M117B had a higher response amplitude in this part of the auditory cortex. **f**, The 2D colormap for the response maps shown in **b**, **c**, and **d**.

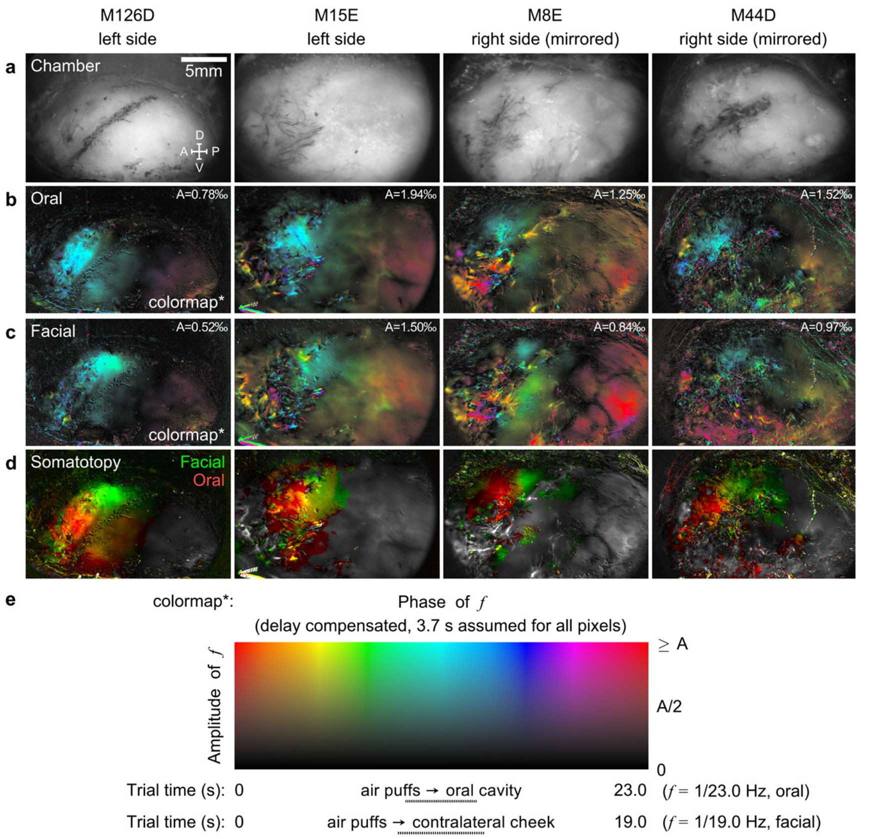


**Extended Data Figure 4.**

**Somatotopy mapping in four tested subjects.** The somatotopy mapping experiment presented in Fig. 2 was performed in four subjects which are shown here in four columns. Subject ID and the imaged hemisphere are indicated on the top of each column. The right hemisphere view is mirrored to match the view of the left hemisphere for display purposes. **a**, The image of the recording chamber in each subject. **b**, The response map evoked by the oral stimuli in each subject. **c**, The response map evoked by the facial stimuli in each subject. A fixed hemodynamic delay value of 3.7 seconds is assumed and compensated for all pixels in these maps (**b**, **c**). Response phase and amplitude are visualized separately as color and intensity in the map, according to the 2D colormap in **e**. For each response map, the upper display limit of amplitude "A" is listed at the top right. **d**, A summary map for each subject, with the oral response on the red color channel, and the facial response on the green color channel. Every subject showed an orofacial somatotopic gradient with the oral component at the more anteroventral side, and the major facial component at the more posterodorsal side. **e,** The 2D colormap for the response maps shown in **b** and **c**.

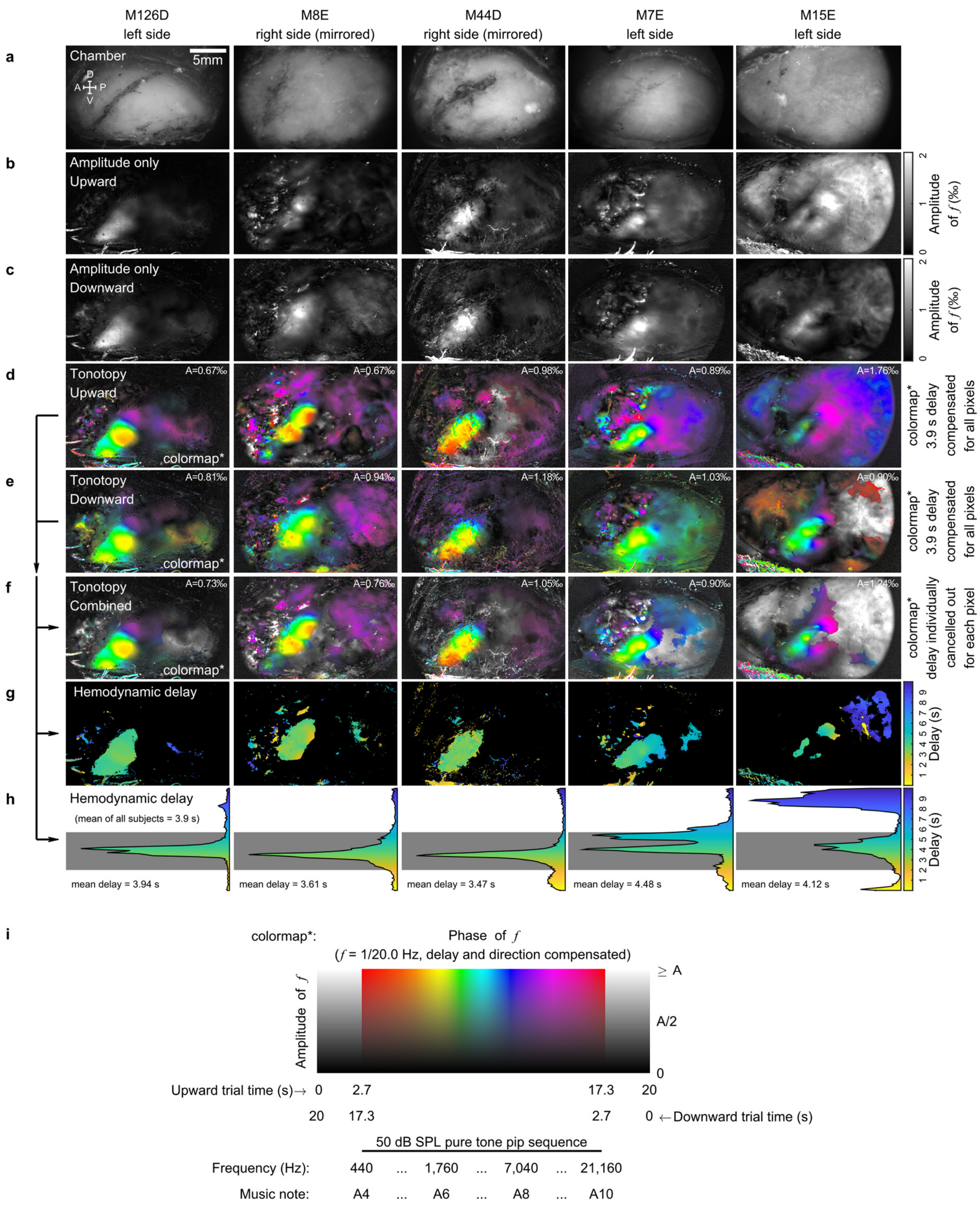


**Extended Data Figure 5.**

**Tonotopy mapping in five tested subjects.** The tonotopy mapping experiment presented in Fig. 2 was performed in five subjects which are shown here in five columns. Subject ID and the imaged hemisphere are indicated on the top of each column. The right hemisphere view is mirrored to match the view of the left hemisphere for display purposes. **a**, The image of the recording chamber in each subject. **b**, Amplitude-only response map of the session with an upward frequency sequence (Fig. 2i right). **c**, Amplitude-only response map of the session with a downward frequency sequence (Fig. 2i left). The region that is consistent with the location of the auditory cortex (Fig. 1b) exhibits the highest response amplitude in most of these maps. The two sessions (**b**, **c**) performed in each subject generally produce very similar maps. **d**, Tonotopic map derived from the session with an upward frequency sequence. **e**, Tonotopic map derived from the session with a downward frequency sequence. The tuning phase and amplitude are visualized separately as color and intensity in these maps (**d**, **e**), according to the 2D colormap in **i**. A fixed hemodynamic delay value of 3.9 seconds is assumed and compensated for all pixels to derive the tuning phase from the raw response phase. **f**, Tonotopic map derived by combing these two "temporally reversed" sessions together. The hemodynamic delay of each pixel was individually estimated and canceled out in this map. **g**, Hemodynamic delay map derived by combing these two "temporally reversed" sessions together. Each hemodynamic delay map only includes the pixels with response amplitudes (average of sessions in **d** and **e**) that are higher than the 90th percentile of all pixels. **h**, Histogram of hemodynamic delay values of the pixels in **g**. Background rectangular gray shade: the acceptable delay range. Bottom number label: averaged delay within this range. The overall mean value of all subjects is 3.9 seconds. For each tonotopic map, the upper display limit of amplitude "A" is listed at the top right. **i**, The 2D colormap for the response maps shown in **d**, **e**, and **f**.

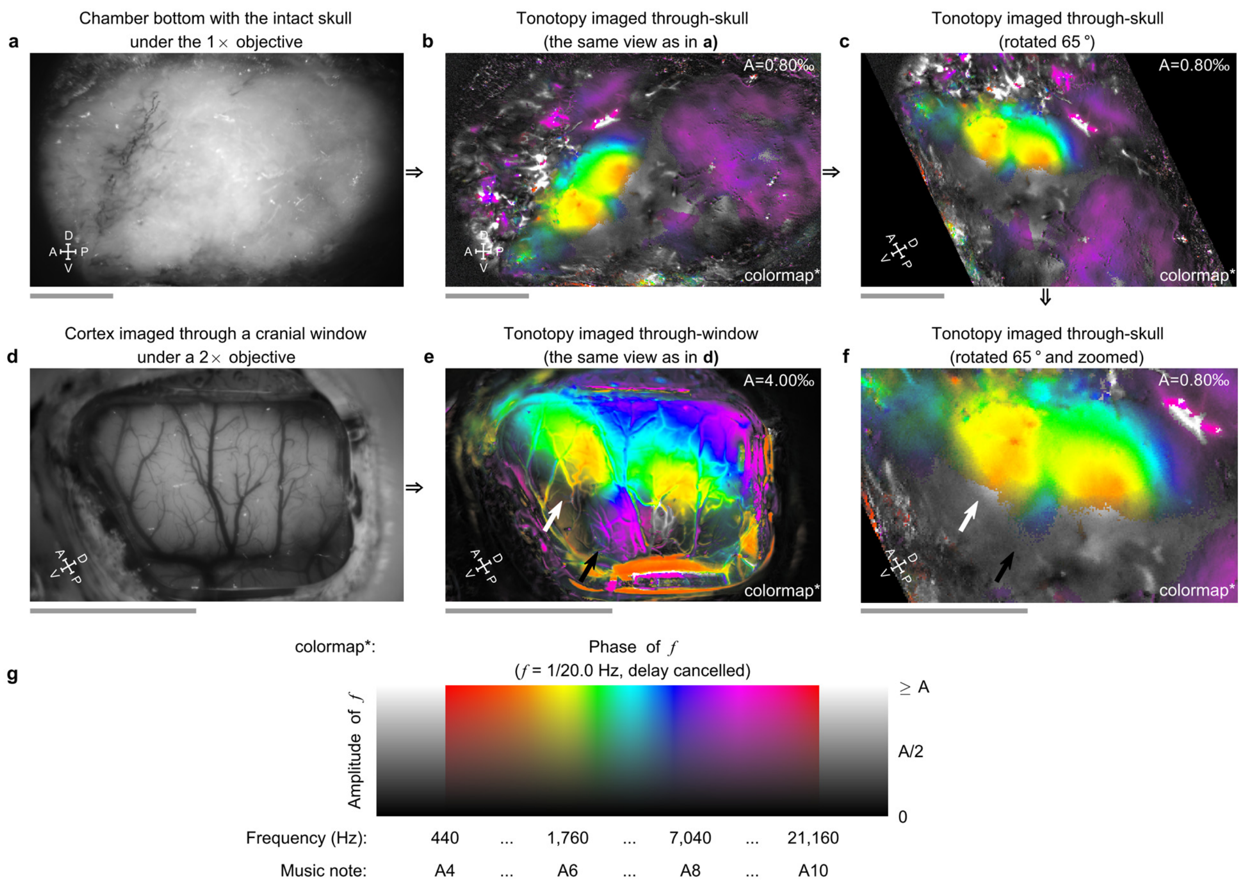


**Extended Data Figure 6.**

**Comparison of tonotopic maps imaged through the intact skull or a cranial window. a**, A view of the recording chamber over the intact skull in subject M8E, imaged with a 1× objective. **b**, The tonotopic map imaged through-skull (the same view as in **a**). **c**, The same map as in **b** but rotated 65° clockwise to position the putative lateral sulcus approximately horizontal. **d**, After the through-skull imaging, subject M8E was chronically implanted with a cranial window over the location of the tonotopic gradients seen in **c** (see Supplementary Information). The cortex was then imaged through the cranial window with a 2× objective. Both the lateral sulcus and the superior temporal sulcus can be seen through the cranial window and match their estimated positions through-skull. **e**, The tonotopic map imaged through the cranial window (the same view as in **d**). A white arrow indicates the location of the low-frequency end of the newly discovered tonotopic gradient (presumably in the RT), whereas a black arrow indicates the location of the high-frequency end of the gradient (presumably in the parabelt). **f**, An enlarged view of the through-skull tonotopic map in **c** to match the through-window view in **e**. The same white and black arrows indicate the low-frequency and high-frequency ends of the newly discovered tonotopic gradient. The through-skull response pattern (**f**) largely matches the through-window response pattern (**e**), suggesting the through-skull imaging indeed measures the cortical intrinsic optical signal, with additional diffusion largely attributed to the skull. The scale of this diffusion is generally consistent with the simulation results (Extended Data Fig. 1e). For each tonotopic map (**b**, **c**, **e**, and **f**), the upper display limit of amplitude "A" is listed at the top right. A=0.80‰ is set for the through-skull maps (**b**, **c**, and **f**), whereas A=4.00‰ is set for the through-window map (**e**). **g**, The 2D colormap for the response maps shown in **b**, **c**, **e**, and **f**. The gray scalebar under each plot in **a-f** indicates a length of 5 mm. A (anterior), P (posterior), D (dorsal), V (ventral). Imaging was performed in the right hemisphere of the subject and the results were mirrored to the left for display purposes.

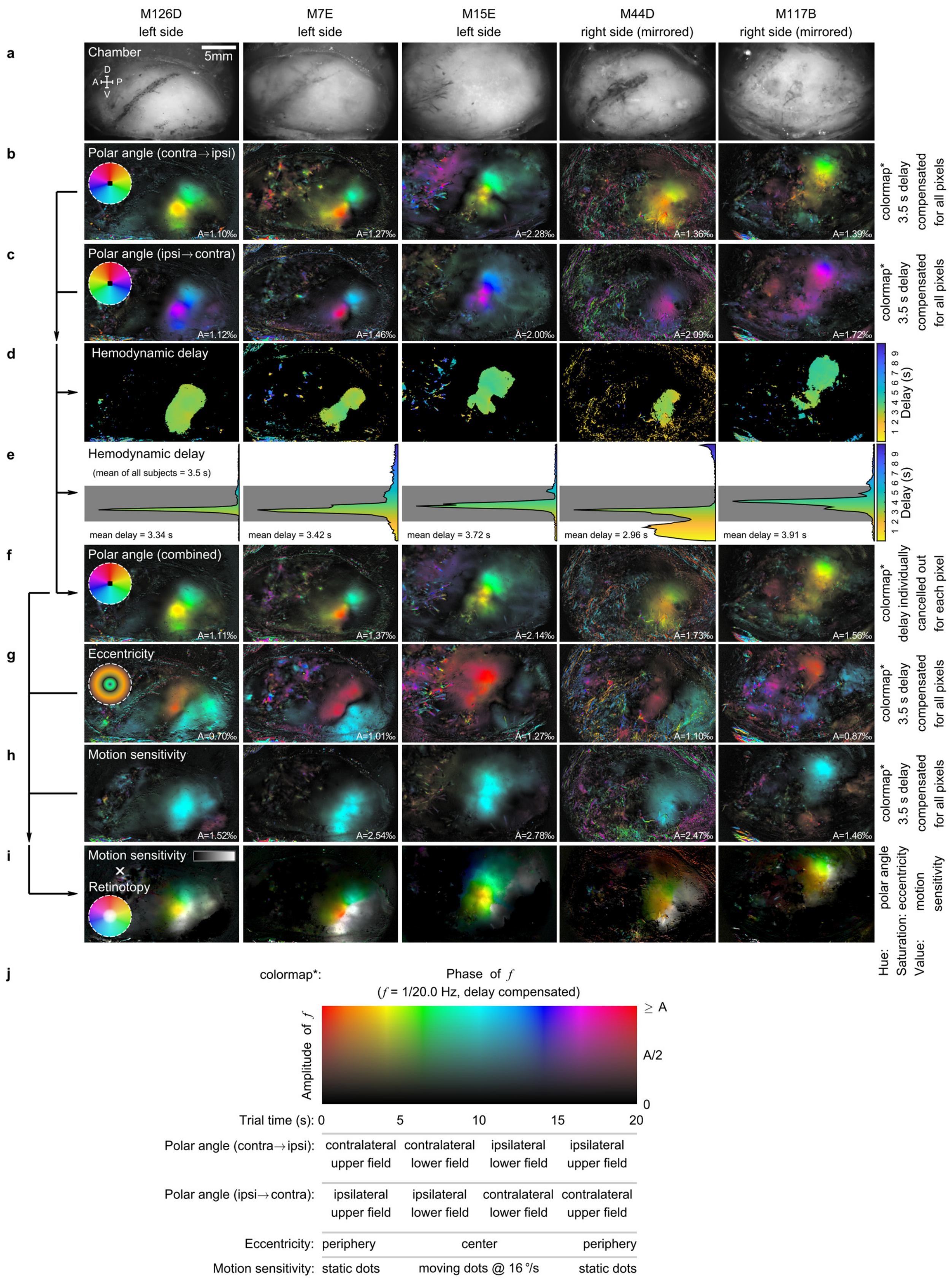

M126D
left side
M7E
left side
M15E
left side
M44D
right side (mirrored)
M117B
right side (mirrored)
a
Chamber
5mm
D
A
P
V
b
Polar angle (contra→ipsi)
A=1.10‰
A=1.27‰
A=2.28‰
A=1.36‰
A=1.39‰
colormap* 3.5 s delay compensated for all pixels
c
Polar angle (ipsi→contra)
A=1.12‰
A=1.46‰
A=2.00‰
A=2.09‰
A=1.72‰
colormap* 3.5 s delay compensated for all pixels
d
Hemodynamic delay
Delay (s)
1 2 3 4 5 6 7 8 9
e
Hemodynamic delay
(mean of all subjects = 3.5 s)
mean delay = 3.34 s
mean delay = 3.42 s
mean delay = 3.72 s
mean delay = 2.96 s
mean delay = 3.91 s
Delay (s)
1 2 3 4 5 6 7 8 9
f
Polar angle (combined)
A=1.11‰
A=1.37‰
A=2.14‰
A=1.73‰
A=1.56‰
colormap* delay individually cancelled out for each pixel
g
Eccentricity
A=0.70‰
A=1.01‰
A=1.27‰
A=1.10‰
A=0.87‰
colormap* 3.5 s delay compensated for all pixels
h
Motion sensitivity
A=1.52‰
A=2.54‰
A=2.78‰
A=2.47‰
A=1.46‰
colormap* 3.5 s delay compensated for all pixels
i
Motion sensitivity
×
Retinotopy
Hue: polar angle
Saturation: eccentricity
Value: motion sensitivity
j
colormap*:
Phase of f
(f = 1/20.0 Hz, delay compensated)
Amplitude of f
≥ A
A/2
0
Trial time (s): 0 5 10 15 20
Polar angle (contra→ipsi): contralateral upper field | contralateral lower field | ipsilateral lower field | ipsilateral upper field
Polar angle (ipsi→contra): ipsilateral upper field | ipsilateral lower field | contralateral lower field | contralateral upper field
Eccentricity: periphery | center | periphery
Motion sensitivity: static dots | moving dots @ 16 °/s | static dots

**Extended Data Figure 7.**

**Retinotopy and motion sensitivity mapping in five tested subjects.** The retinotopy and motion sensitivity mapping experiments presented in Fig. 3 were performed in five subjects which are shown here in five columns. Subject ID and the imaged hemisphere are indicated on the top of each column. The right hemisphere view is mirrored to match the view of the left hemisphere for display purposes. **a**, The image of the recording chamber in each subject. **b**, The retinotopic polar angle map, derived from the session with the motion range sweeping clockwise for left hemispheres, or counterclockwise for right hemispheres (thus, relative to the imaging side, each cycle had the visual field swept in a sequence: contralateral-upper, contralateral-lower, ipsilateral-lower, and ipsilateral-upper). **c**, The retinotopic polar angle map, derived from the session with the motion range sweeping counterclockwise for left hemispheres, or clockwise for right hemispheres (thus, relative to the imaging side, each cycle had the visual field swept in a sequence: ipsilateral-upper, ipsilateral-lower, contralateral-lower, and contralateral-upper). A fixed hemodynamic delay value of 3.5 seconds is assumed and compensated for all pixels to derive the tuning phase from the raw response phase. The tuning phase and amplitude are visualized separately as color and intensity in the map, according to the 2D colormap in **j**. The colormap is transformed into a visual field color code and shown as a color wheel at the upper left (coded for left hemispheres). The upper display limit of amplitude “A” is labeled at the bottom right in each map. **d**, Hemodynamic delay map derived by combing these two “temporally reversed” sessions together. Each hemodynamic delay map only includes the pixels with response amplitudes (average of the sessions in **b** and **c**) that are higher than the 90th percentile of all pixels. **e**, The histogram of hemodynamic delay values of the pixels in **d**. Background rectangular gray shade: the acceptable delay range. Bottom number label: averaged delay within this range. The overall mean value of all subjects is 3.5 seconds. **f**, Retinotopic polar angle map derived by combing these two "temporally reversed" sessions (**b**, **c**) together. The hemodynamic delay of each pixel is individually estimated and canceled out in this map. **g**, The retinotopic eccentricity map. **h**, The motion sensitivity map. Each of these maps is derived from one of the other two independent recording sessions. A fixed hemodynamic delay value of 3.5 seconds is assumed and compensated for all pixels to derive the tuning phase from the raw response phase. The tuning phase and amplitude are visualized separately as color and intensity in the map, according to the 2D colormap in **j**. The colormap is transformed into a visual field color code for the eccentricity session and shown as a color wheel at the upper left. The upper display limit of amplitude "A" is labeled at the bottom right in each map. **i**, The summary map by combining the retinotopic maps and the motion sensitivity map. These maps are encoded into separate channels in the HSV color space. The retinotopic polar angle tuning (**f**) was encoded in the hue (color) channel, whereas the retinotopic eccentricity tuning (**g**) was encoded in the saturation (chroma) channel. A visual field color wheel incorporating these 2 features is shown at the lower left (for left hemispheres). A pixel sensitive to the visual field center (<5° radius) would be shown in white, whereas a pixel sensitive to the periphery (>5° radius) would be shown in a vivid color according to its polar angle tuning. Meanwhile, the motion sensitivity (**h**) is encoded in the value (brightness) channel. The more motion sensitivity a pixel exhibited to moving dots, the brighter it appears in the map. Together, it is evident that the motion sensitivity to moving dots extends across two retinotopically organized regions. Each region has a representation of the contralateral visual field. While the eccentricity gradient is continuous across these two regions, the polar angle gradient reverses at the boundary between them. **j,** The 2D colormap for the response maps shown in **b**, **c**, **f**, **g**, and **h**.

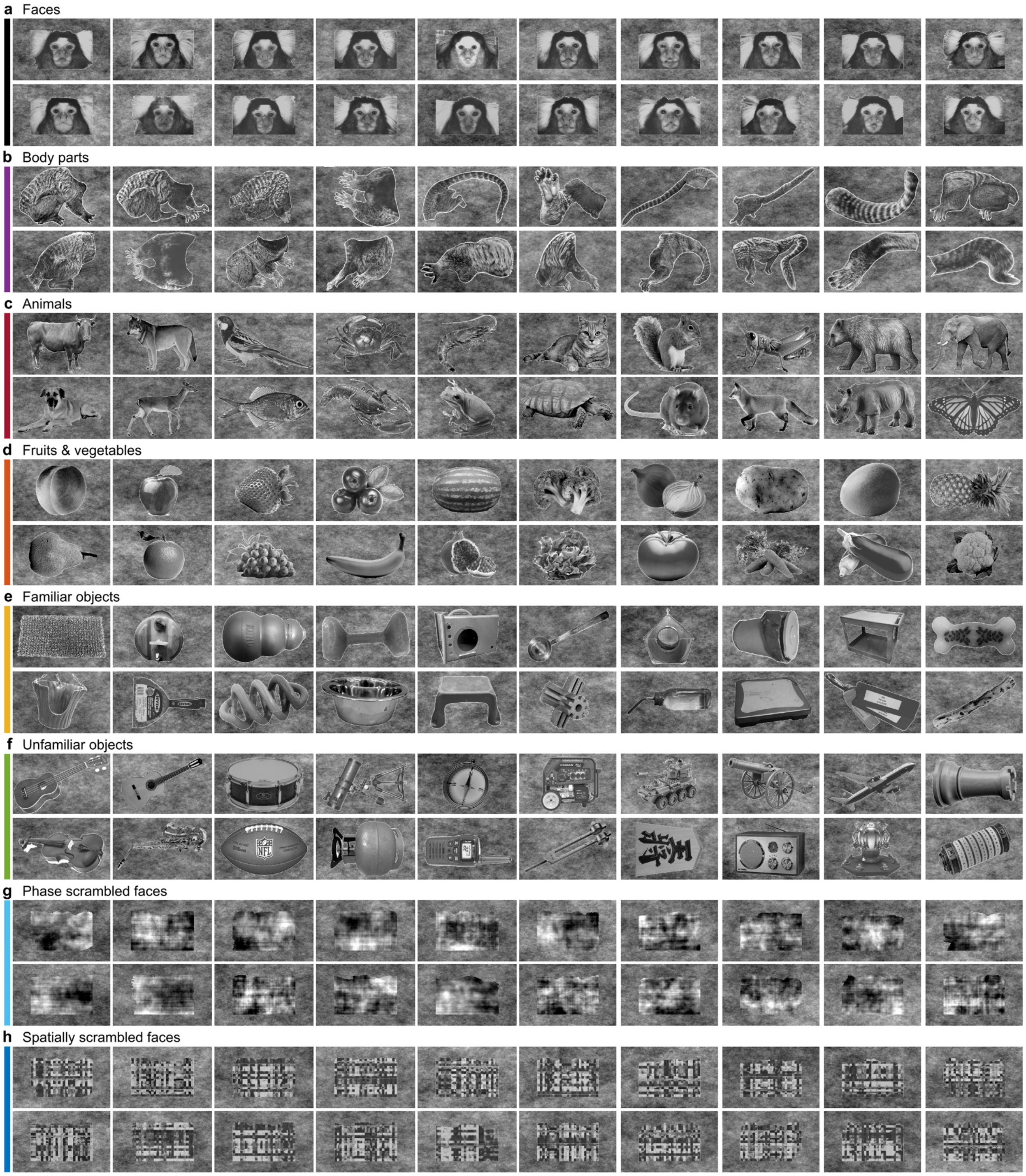


**Extended Data Figure 8.**

**Face patch mapping stimuli.** Eight visual form categories were used in the face patch mapping experiment, including faces (**a**), body parts (**b**), animals (**c**), fruits and vegetables (**d**), familiar objects (**e**), unfamiliar objects (**f**), phase scrambled faces (**g**), and spatially scrambled faces (**h**). Each category had 20 exemplars. Together, 160 exemplars used are all shown. The marmoset faces were used in the study by Hung et. al.[44] and provided to us as a courtesy by the authors of that study.

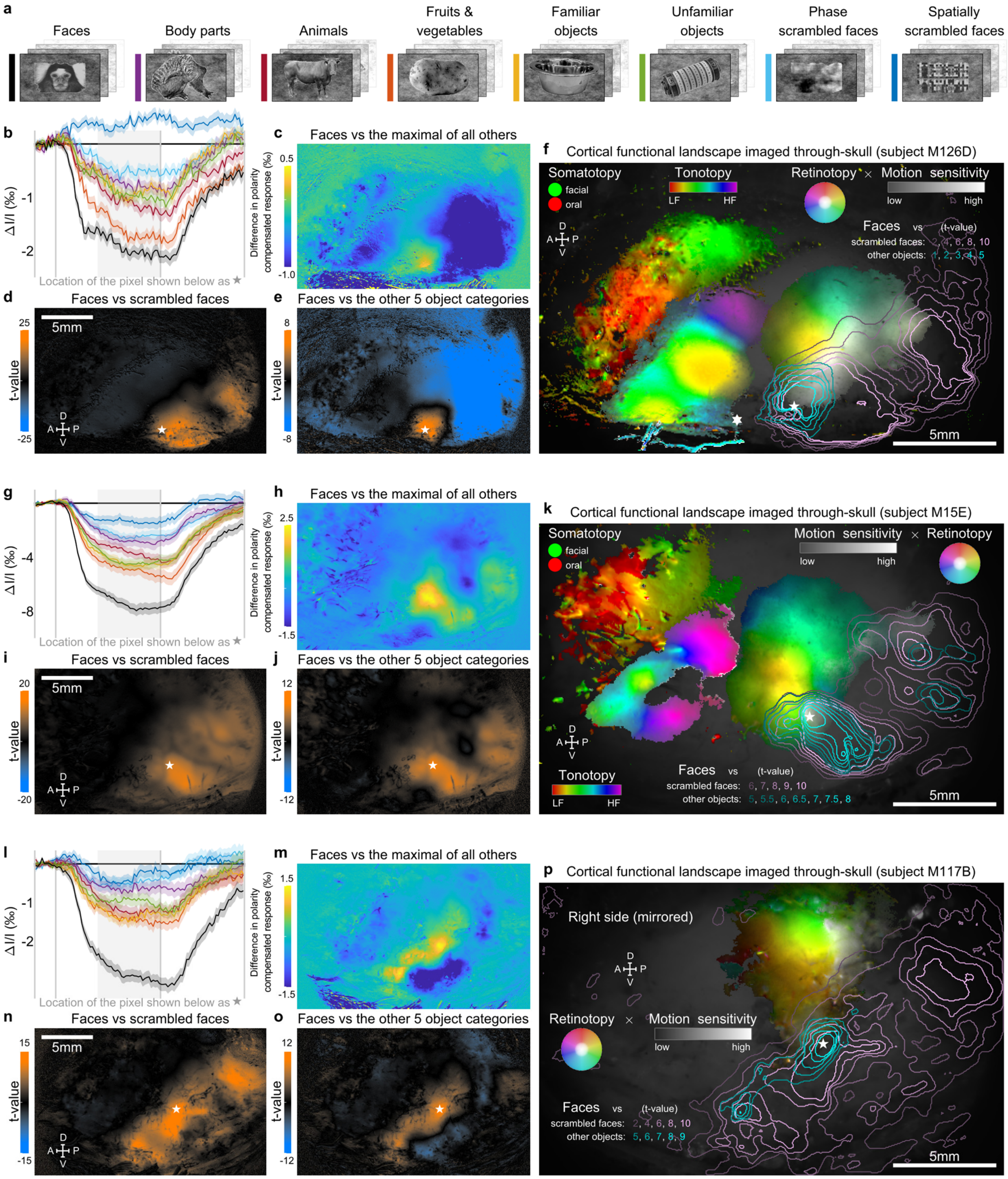


1

2 **Extended Data Figure 9.**

**Face patch mapping and cortical functional landscapes assembled from all through-skull maps in three tested subjects.** The face patch mapping experiment (Fig. 4) was performed in three subjects: M126D, M15E, and M117B. **a**, Eight categories of visual stimuli, with their exemplars and color code (used in **b**, **g**, and **l** as well). **b-f**, Plots for subject M126D, as appeared in Fig. 4d-h, re-listed here for the comparison purpose (n=218 for each category). **g**, Response to each category of a face-sensitive pixel in subject M15E (location labeled by a pentagram in both **i** and **j**). The colors indicate categories, as shown in **a**. Each average response trace is shown as a solid curve with a shade representing the corresponding SEM (n=134). The responses to the faces are the strongest among all categories, followed by the responses to other object categories and then the scrambled faces. The rectangular background gray shade indicates an averaging time window for calculating a response value of each trial, with signal polarity compensated. **h**, Differential response map between the mean response to the face category and the maximum of mean responses to all other categories (n=134 for each category). A positive value in the map indicates that pixel responded more strongly to faces than to any other 7 stimulus categories. **i**. The t-value map for comparing faces versus scrambled faces. A t-value is calculated for each pixel in the map by comparing its single-trial responses between those to the faces (n=134) and those to the scrambled faces (n=268). Scrambling sensitivity in the map (orange color: high t-values) implicates involvement in visual form processing[49]. **j**. The t-value map for comparing the face category (n=134) versus all other 5 object categories (n=670). The face-over-object sensitivity in the map (orange color: high t-values) defines the location of a face patch. **k**. The cortical functional landscape summarized from modality-specific maps and overlaid on a dimmed image of the recording chamber. The motion-sensitivity is shown with the retinotopic tunings together in the "HSV" color space as in Extended Data Fig. 7i (hue [color]: polar angle;

saturation [chroma]: eccentricity; value [brightness]: motion sensitivity). The t-value maps are shown by the iso-t-value contour lines. Pentagram: a face-sensitive pixel (same location as in **i** and **j**). The face patch (outlined by the blue contours) is largely overlapped with both the moving-dots-sensitive region and the retinotopic region representing the lower center of the visual field. **l**, Response to each category of a face-sensitive pixel in subject M117B (n=234, location labeled by a pentagram in both **n** and **o**). The panel is plotted in the same way as in **g**. The responses to the faces are the strongest among all categories, followed by the responses to other object categories and then the scrambled faces. **m**, Differential response map between the mean response to the face category and the maximum of mean responses to all other categories (n=234 for each category). Since imaging was performed in the right hemisphere of subject M117B, all maps and the visual field color code of this subject are mirrored to match the other subjects for display purposes. **n**. The t-value map for comparing faces (n=234) versus scrambled faces (n=468). **o**. The t-value map for comparing the face category (n=234) versus all other 5 object categories (n=1170). The face-over-object sensitivity in the map (orange color: high t-values) defines the location of a face patch. **p**. The cortical functional landscape summarized from the visual maps and overlaid on a dimmed image of the recording chamber. The motion-sensitivity is shown with the retinotopic tunings together in the "HSV" color space as in Extended Data Fig. 7i (hue [color]: polar angle; saturation [chroma]: eccentricity; value [brightness]: motion sensitivity). The t-value maps are shown by the iso-t-value contour lines. Pentagram: a face-sensitive pixel (same location as in **n** and **o**). The face patch (outlined by the blue contours) is largely overlapped with both the moving-dots-sensitive region and the retinotopic region representing the lower center of the visual field.

### EXTENDED DATA TABLES

| Subject ID | M126D | M15E | M8E | M7E | M44D | M117B |
|---|---|---|---|---|---|---|
| Gender | female | male | male | male | male | male |
| Imaging side | left | left | right | left | right | right |
| Chamber bottom material | orthodontic resin | C&B metabond | orthodontic resin | orthodontic resin | orthodontic resin | C&B metabond |
| Age at the head cap implantation (month) | 36 | 43 | 45 | 44 | 38 | 54 |
| Age at the earliest imaging acquisition (month) | 47 | 44 | 46 | 46 | 54 | 80 |
| Age at the latest imaging acquisition (month) | 52 | 48 | 48 | 46 | 55 | 81 |
| Experiment participation: modality parcellation | Y | Y | Y | Y | Y | Y |
| Experiment participation: somatotopy | Y | Y | Y | N/A[1] | Y | N/A[2] |
| Experiment participation: tonotopy | Y | Y | Y | Y | Y | N/A[3] |
| Experiment participation: retinotopy | Y | Y | N/A[4] | Y | Y | Y |
| Experiment participation: face patch | Y | Y | N/A[4] | N/A[1] | N/A[5] | Y |
| Availability comments | [1] The subject was not available anymore when these experiments were designed.<br>[2] The parcellation experiment showed the recording chamber of this subject included very little somatosensory cortex for further investigations.<br>[3] The skull over part of the auditory cortex in this subject was thinned in an earlier pilot experiment. Thus tonotopy mapping through the intact unthinned skull could not be performed anymore in this subject<br>[4] Pursuit eye movement behavior in this subject was observed abnormal comparing to that in other subjects (confirmed by eye tracking). Thus the subject was excluded from further visual experiments<br>[5] The posteroventral part of the chamber was not big enough to include the full region of interest in this subject, based on the results of the retinotopy mapping experiment. Thus further face patch searching was not performed | | | | | |

**Extended Data Table 1.**

**Summary of tested subjects.** A breakdown summary of all six tested subjects in the current study, including one female, five males; three left hemispheres, three right hemispheres; four with chamber bottom covered with orthodontic resin, two with chamber bottom covered with C&B Metabond. Subjects' ages ranged from 36 to 54 months old at the time of head cap implantation, and from 44 to 81 months old during imaging acquisition. Out of 30 possible experiments (5 experiments × 6 subjects), 7 were not conducted due to the availability issues commented on in the table.
