## Supplementary Information for "Functional maps of the primate cortex revealed by through-skull wide-field optical imaging"

### Optical properties of the skull and the brain.

To analyze how the imaging light propagates in the tissue (Extended Data Fig. 1), the optical properties of both the skull and the brain were obtained from previous studies. For scattering in the skull, we followed the equation as $\mu'_s(\lambda) = 153.302 \cdot \lambda^{-0.65} (mm^{-1})$, where $\lambda$ is the wavelength in nm, and $\mu'_s(\lambda)$ is the reduced scattering coefficient[66]. For absorption in the skull, coefficients were averaged across three datasets to cover the spectrum[67-69]. For scattering in the brain, the scattering coefficients $\mu_s(\lambda)$ and the anisotropy factor (the mean cosine of the scattering angle) $g$ were obtained across a wide spectrum[69]. $\mu'_s(\lambda)$ were calculated as $\mu'_s(\lambda) = \mu_s(\lambda) \cdot (1-g)$. For absorption in the brain, we assumed: (i) the major absorber in the brain is blood; (ii) blood composes 3% volume of the brain; (iii) the concentration of hemoglobin in blood is 2mM (~130g/L); and (iv) the composition of hemoglobin is 25% HbR and 75% HbO. The absorption coefficients of the brain were thus derived based on the absorption spectra of HbR and HbO (https://omlc.org/spectra/hemoglobin/summary.html). These coefficients are plotted in Extended Data Fig. 1a and b.

### Estimating the photon visitation depth in the skull and the brain.

A previous study simulated and analyzed the photon visitation depth of polarized light backscattered from a scattering media[21]. A semi-infinite homogeneous medium was assumed to be absorption-free and comprised of Mie scatterers with the anisotropy factor $g$ of 0.92 and the scatterer size of 2.21 wavelengths. Illumination was simulated as a linearly polarized beam placed perpendicular to the medium surface. Photons were collected either through the co-linearly polarized channel or through the cross-linearly polarized channel (polarization direction is relative to the polarization of the illumination). Photons that undergo relatively few scattering events and maintain their initial polarization contribute only to the co-linear channel, whereas photons that have random polarization through multiple scattering events contribute equally to the co-linear channel and cross-linear channel. The "linear-maintaining" channel was thus defined as the intensity difference between the "co-linear" channel and the "cross-linear" channel. We borrowed the estimated visitation depths of the "linear-maintaining" and the "cross-linear" channels from this study[21]. These visitation depths were expressed in the unit of the mean free path (MFP). To translate these results into a model consists of two layers (the skull on top of the brain) with absolute thicknesses, we calculated MFPs [ $MFP = 1/\mu_s(\lambda)$ ] for both the skull and the brain, based on the parameters mentioned above (absolute visitation depths shown in Extended Data Fig. 1d). It is worth noting, there are several assumptions in the current model that may affect the degree it resembles the real condition: (i) the refractive index of the skull and the brain were assumed equal (a homogeneous medium was artificially cut into two layers); (ii) both layers were assumed absorption-free; and (iii) no other structures were assumed between the skull and the brain, such as the dura mater. All these factors may contribute to a bias in estimating an absolute visitation depth. Nevertheless, the relative scale of the visitation depths estimated here may provide a reference to guide practice.

**Estimating the lateral resolution of XINTRINSIC.**

We used Monte Carlo (MC) simulation to estimate the lateral resolution of our imaging system. In the simulation model, the medium consisted of 3 stacking layers, each mimicking the skull, the gray matter, and the white matter. For the backscattered light collected from a point on the surface of the skull, its intensity would be affected if there is an absorption change in the gray matter underneath the skull. The more laterally displaced the change is from the point, the less sensitive the response is to the change. The lateral resolution of the signal recorded on this point was described as the normalized lateral profile of its sensitivity to the absorption change in the gray matter. We did not differentiate photons by their polarization, since the assumed thickness of the skull, 0.50 mm [(6)], is thicker than the travel distance needed for linearly polarized light to be completely depolarized[70] ($1/\mu'_s(\lambda)$, estimated as 0.47 mm for green light). For photons that eventually contribute to the sensitivity of the absorption change, they must travel through the skull to reach the gray matter and are thus already depolarized. Therefore, differentiating these photons by their polarization states would not result in much difference in estimating the lateral resolution here.

The reciprocity rule is used in our simulation, that is, the trajectory of light propagation in one direction is equivalent to the reversed direction. To simulate collecting photons that exit the medium from a specific point on the skull surface, we reversely launched photons into the medium at this point. The launched photons may eventually leave the medium from the top surface and be collected, which reversely simulates the illumination process. In our real experiments, the epi-illumination was applied through the same objective (NA=0.03) that collected backscattered light (the back focal aperture of the objective was fulfilled in illumination). We thus limited the photon launching and collection angles in the simulation within this range (NA≤0.03).

The simulation was performed in Matlab (Mathworks) with a GPU-based simulation package[71] (MCXLAB). A medium was set to be a finite cube with a dimension of 10×10×10 $mm^3$. The three layers of the medium (skull, gray matter, and white matter) were configured according to the parameters listed in Supplementary Table 1. In the initial photon trajectory simulation, all layers of the medium were first set free of any absorption. $1\times10^9$ photons were launched into the medium at the center point of the surface. The simulated trajectory of each launched photon was collected if its exiting angle from the medium is within the collection angle range (NA≤0.03).

| | Wavelength (nm) | 470 | 530 | 590 | 625 | 730 | 850 |
|---|---|---|---|---|---|---|---|
| Skull: $n$ = 1.56, thickness = 0.5 mm | $\mu_s(\lambda)(mm^{-1})$ | 28.10 | 25.99 | 24.24 | 23.35 | 21.11 | 19.12 |
| | $\mu_a(\lambda)(mm^{-1})$ | 0.1137 | 0.1137 | 0.0750 | 0.0424 | 0.0274 | 0.0172 |
| | $g$ | 0.920 | 0.920 | 0.920 | 0.920 | 0.920 | 0.920 |
| Gray matter: $n$ = 1.37, thickness = 1.3 mm | $\mu_s(\lambda)(mm^{-1})$ | 11.63 | 10.53 | 9.62 | 9.15 | 8.12 | 7.24 |
| | $\mu_a(\lambda)(mm^{-1})$ | 0.465 | 0.638 | 0.287 | 0.032 | 0.009 | 0.016 |
| | $g$ | 0.882 | 0.887 | 0.892 | 0.895 | 0.899 | 0.898 |
| White matter: $n$ = 1.37 thickness = 8.2 mm | $\mu_s(\lambda)(mm^{-1})$ | 42.85 | 41.94 | 40.70 | 40.28 | 38.61 | 35.31 |
| | $\mu_a(\lambda)(mm^{-1})$ | 0.465 | 0.638 | 0.287 | 0.032 | 0.009 | 0.016 |
| | $g$ | 0.790 | 0.810 | 0.830 | 0.836 | 0.858 | 0.871 |

**Supplementary Table 1.**

**Optical and physical properties of the skull and the brain.** Optical properties of the skull and the brain were derived from the same data sources and assumptions mentioned in the previous section. An anisotropy factor $g$ = 0.92 was further assumed for the skull[21]. Other values, such as skull thickness[6], gray matter thickness[73], and refractive indices ($n$) of the skull[74] and the brain[71] were also listed.

To put absorption back into consideration, the weight ($W$) of each photon was determined by the Beer's Law based on its simulated trajectory:

$$W = e^{-\sum_{i=1}^{n} \mu_a(i) \cdot x(i)}$$

Where $x(i)$ is the pathlength of a photon traveled in the $i$th medium, and $\mu_a(i)$ is the absorption coefficient of that medium. Therefore, the sum of these detected photons' weights would reflect the light intensity detected by the camera.

To estimate the system's lateral resolution defined above, we divided the gray matter layer into a 2D grid of column-shaped voxels, each had a size of 0.033 mm × 0.033 mm × 1.300 mm (lateral width, lateral width, depth). To apply a perturbation MC method[72], we increased the absorption coefficient by 10% in a single gray matter voxel and recalculated each photon's weight based on its originally simulated trajectory. The resulting change in the total light intensity (ΔI) thus reflects the imaging point's sensitivity to the voxel's absorption change. The same process was repeated for the center row of voxels that passes the imaging point. A normalized ΔI curve was plotted against the lateral distance of these voxels from the imaging point (Extended Data Fig. 1e). This curve reflects the lateral profile of the imaging point's sensitivity to the absorption changes in these gray matter voxels. We used HWHM (half-width at half maximum) and standard deviation (σ) of the lateral profile as measurements of the lateral resolution.

**Intrinsic optical signal imaging through a chronically implanted cranial window.**

The basic procedures for chronically implanting an artificial dura based cranial window in NHPs have been previously described[5]. After through-skull imaging, the mapping results were used to mark a target position for a subsequent craniotomy within the imaging chamber. To protect the planned craniotomy, a rubber piston cover was taken from the plunger of a standard 10 ml disposable syringe and put on top of the targeted position. Additional dental cement material was added for ~1mm in height around the rubber cover to form a socket and prevent the cover from sliding. The window implantation surgery was performed under sterile conditions when the animal was anesthetized by isoflurane (0.5-2.0%, mixed with pure oxygen). The rubber cover was taken away and the craniotomy and durotomy over the planned position were performed. An artificial dura pre-molded with silicone in a hat-like shape[5] was then implanted. The gap between the craniotomy edge and the sidewall of the artificial dura was filled and sealed with surgical silicone adhesive (WPI, Kwik-Sil). The rubber cover was put back into the pre-built socket to protect the cranial window and sealed to the original recording chamber wall by a dental impression material (GC America, EXAMIX NDS injection). All experimental procedures were approved by the Johns Hopkins University Animal Use and Care Committee.

After the animal fully recovered from the window implantation surgery, intrinsic optical signal imaging was performed through the cranial window (Extended Data Fig. 6d). Since the cranial window is ~9.5 mm × 6.4 mm in size (with one corner retracted) to target the entire auditory cortex on the brain surface, we swapped out the 1× objective (used for all through-skull experiments) and replaced it with a 2× objective (Thorlabs, TL2X-SAP). The same tonotopy mapping experiment (Fig. 2i) was performed through the cranial window, this time with 100 cycles instead of 20 cycles for both the “upward” and the “downward” sessions. The results were analyzed and visualized (Extended Data Fig. 6e) following the same way as in through-skull analysis.
